# Decoding the transcriptome of atherosclerotic plaque at single-cell resolution

**DOI:** 10.1101/2020.03.03.968123

**Authors:** Tom Alsaigh, Doug Evans, David Frankel, Ali Torkamani

## Abstract

Atherogenesis involves an interplay of inflammation, tissue remodeling and cellular transdifferentiation (CTD), making it especially difficult to precisely delineate its pathophysiology. Here we examine the single-cell transcriptome of entire atherosclerotic core (AC) plaques and patient-matched proximal adjacent (PA) portions of carotid artery tissue from patients undergoing carotid endarterectomy. We use a novel tissue dissociation strategy, single-cell RNA sequencing, and systems-biology approaches to analyze the transcriptional profiles of six main cell populations and identify key gene drivers of pathogenic biological processes in vascular smooth muscle cells (VSMCs) and endothelial cells (ECs). Our results reveal an anatomic continuum whereby PA cells promote and respond to inflammatory processes and eventually transition through CTD into matrix-secreting cells in the AC. Inflammatory signaling in PA ECs is driven by IL6, while TNFa signaling defines inflammation in both PA ECs and VSMCs. Furthermore, we identify *POSTN, SPP1 and IBSP* in AC VSMCs, and *ITLN1, SCX* and *S100A4* in AC ECs as key drivers of CTD in the atherosclerotic core. These results establish an anatomic framework for atherogenesis and suggest a site-specific strategy for disruption of disease progression.

## Introduction

The pathophysiology of atherosclerosis is exceptionally complex and involves inflammation^1^, cellular transdifferentiation^2,3^, and dynamic interactions between a variety of cell types within the vascular wall^4,5^. For example, the phenotypic landscape of VSMCs is increasingly understood to be quite dynamic as these cells often participate in phenotype switching^6^ and contribute to the development of extracellular matrix producing cells in atherosclerosis^4^. Many of these pathogenic processes are mediated through molecular crosstalk between VSMCs^7^, however disease progression in general is increasingly appreciated to involve coordinated genomic and molecular communication between the plethora of immune cell subtypes and other cellular components of the arterial wall^8,9^.

Evidence from genome-wide association studies (GWAS) suggests a plurality of genetic drivers of atherosclerosis acting through tissue-specific regulation^10^, underscoring the importance of uncovering cell-type specific genetic drivers of disease in order to expose new therapeutic opportunities. Efforts made using microarrays to evaluate gene expression changes in patients with carotid artery plaque have yielded important information^11^, however, these bulk sequencing studies obscure the diverse plaque environment by assessing complete transcriptome profiles without distinction between predominant cell types contributing to gene expression.

More recent studies have advanced our understanding of the rich cellular heterogeneity within the plaque environment through single-cell transcriptomics approaches. For example, in murine atherosclerotic aortas three distinct macrophage subsets were identified, including inflammatory, Res-like and TREM2hi macrophages^12^, enhancing our understanding of macrophage diversity within plaque. Additionally, immunophenotyping in carotid atherosclerosis revealed fundamental differences between T-cells and macrophages from symptomatic versus asymptomatic patients, including expansion of the CD4+ T-cell subset and macrophages with varied phenotypes in symptomatic patients^13^. However, these studies isolated specific cell lineages and excluded the site of disease – the vascular wall.

The interplay between immune and vascular wall cells participating in diverse inflammatory and remodeling processes in atherosclerosis underscores the advantages of using unbiased single-cell transcriptomics to interrogate major cell types and key genomic drivers of disease progression. Here we developed a tissue dissociation strategy to allow for single-cell RNA sequencing (scRNAseq) of atherosclerotic core (AC) plaques and patient-matched proximal adjacent (PA) portions of the carotid artery in their entirety, without preference for cell type, providing an unbiased view of disease progression. This strategy, in concert with systems biology approaches, has allowed us to identify key transcriptional drivers of disease progression in the vascular wall. Together, this study identifies new therapeutic opportunities and lays the groundwork for future investigation to illuminate new biomarkers and risk stratification tools for clinical application.

## Results

### Tissue source and processing

Paired sections of tissue, including both artery and plaque, were recovered from the atherosclerotic core (AC) and proximally adjacent (PA) region of three patients with type VII calcified plaques undergoing endarterectomy (Fig 1a, Supplementary Table 1). Due to the rich cellular composition of carotid artery and plethora of debris in plaque (ie lipid, fibrinogen, etc), dissociation and generation of single-cell suspensions amenable to single-cell RNA sequencing was difficult. After tissue collection, enzymatic digestion, RBC lysis and filtration were the initial steps required to generate single-cells (see *Methods* and Fig. 1b). However, despite efficient enzymatic dissociation and significant filtering of our sample, we were still challenged by abundant plaque debris, which ultimately resulted in poor single-cell capture rates. In order to overcome this issue without isolating specific cell types through cell-marker antibody labeling, we devised a strategy to label all cells in the sample with a far-red excitation-emission live/dead cell nuclear stain (DRAQ5). All cells in the sample were stained, with debris being left unstained by the dye. Subsequently, DRAQ5+ cells were manually gated and sorted from the remainder of the debris using FACS. Cells isolated from the entire filtered sample represented <1% of the total particles in the sample (Extended Data Fig. 1a-f). Viability of remaining cells was assessed and was always >80% using this technique for cell separation (see *Methods*). The cells were then processed for single-cell sequencing.

**Fig. 1:**
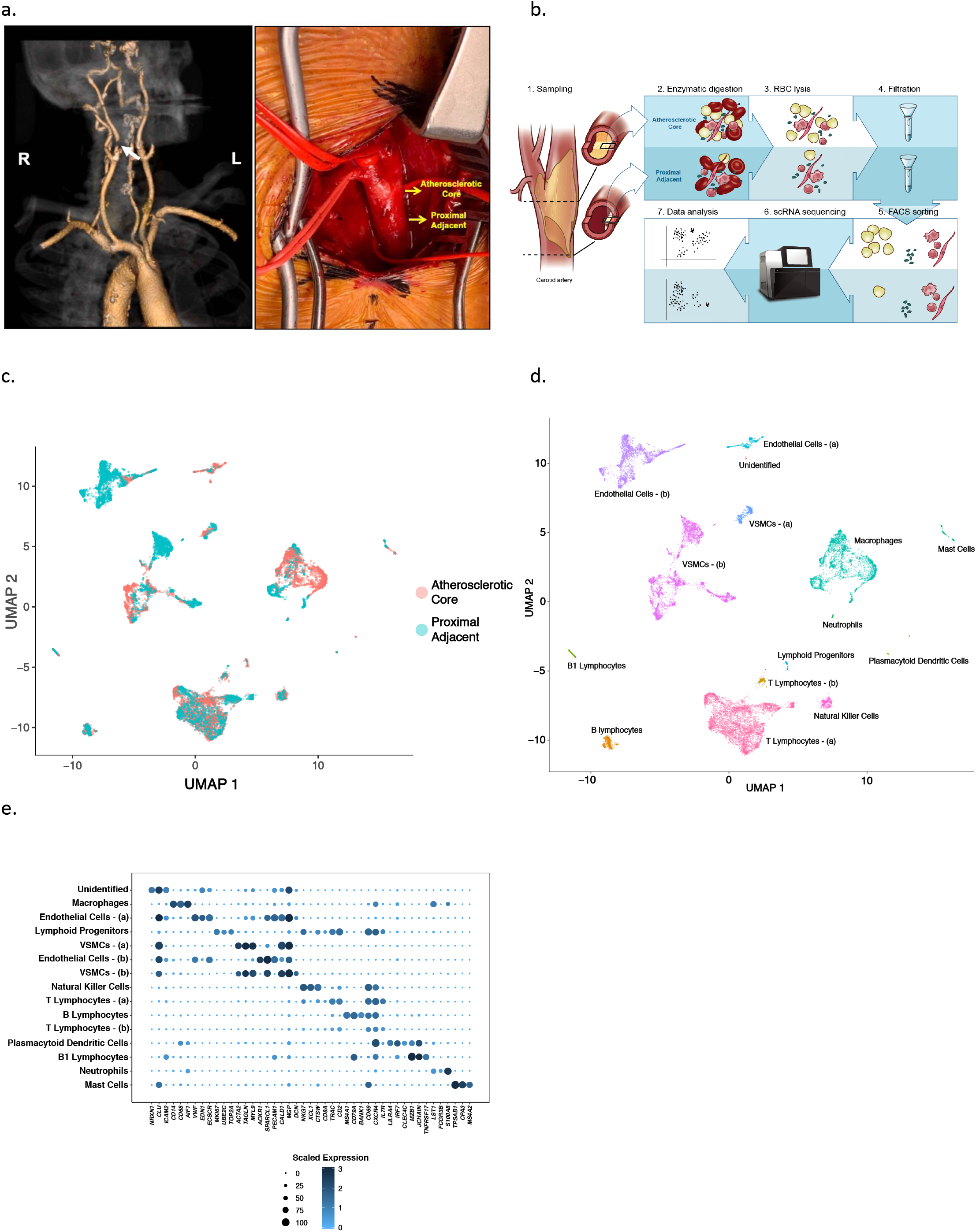
Complete dissociation and scRNAseq profiling of 51,981 cells from the AC and PA region results in 15 distinct cell populations. **a,** Computed tomography angiogram demonstrating right internal carotid occlusion with >70% stenosis due to plaque (left panel); anatomic location of dissected tissue for experiments presented (right panel). **b,** Schematic depicting steps necessary for complete dissociation of artery and plaque from their respective anatomic regions (details in Methods). **c,d** UMAP visualization of down-sampled cells (n=17,100 cells) split by anatomic location **(c)**, and by cell-type **(d)**. **e**, cell-type marker genes for 15 distinct cell populations presented as a dotplot. Dot size depicts the fraction of cells expressing a gene. Dot color depicts the degree of expression of each gene.

### Cell-type identification

Generation of single-cells from three patient-matched AC and PA samples yielded 51,981 cells total, with an average of ~13,000 AC cells/patient and ~5,000 PA cells/patient. Cell number disparities are due to the difference in size of the AC vs PA tissue itself. All datasets were down-sampled randomly so that the cell number per dataset was equal across samples. UMAP-based clustering (see *Methods*) of this down-sampled dataset reveals 15 distinct cell partitions (Fig. 1c,d), representing 17,100 cells total. In order to categorize partitions into major cell types we examined genes expressed in greater than 80% of cells and at a mean expression count greater than 2. A dotplot representing three marker genes hand-selected for each partition is presented in (Fig. 1e). Assigned cell-types in order from greatest (4,944 cells) to least (24 cells) abundant included: T-lymphocytes, macrophages, VSMCs, ECs, B-lymphocytes, natural killer cells, B1-lymphocytes, mast cells, lymphoid progenitors, plasmacytoid dendritic cells, and an unidentified partition (Supplementary Table 2). Following doublet filtering using a marker-gene exclusion method (see *Methods*), and removal of partitions with too few cells for differential gene expression analysis, the remaining cells were re-clustered producing six major partitions: macrophages, ECs, VSMCs, NKT cells, T- and B-lymphocytes (Fig. 2a-c, Extended Data Figs. 2,3). We then assessed differential gene expression between AC and PA regions across these 6 major cell types.

**Fig. 2:**
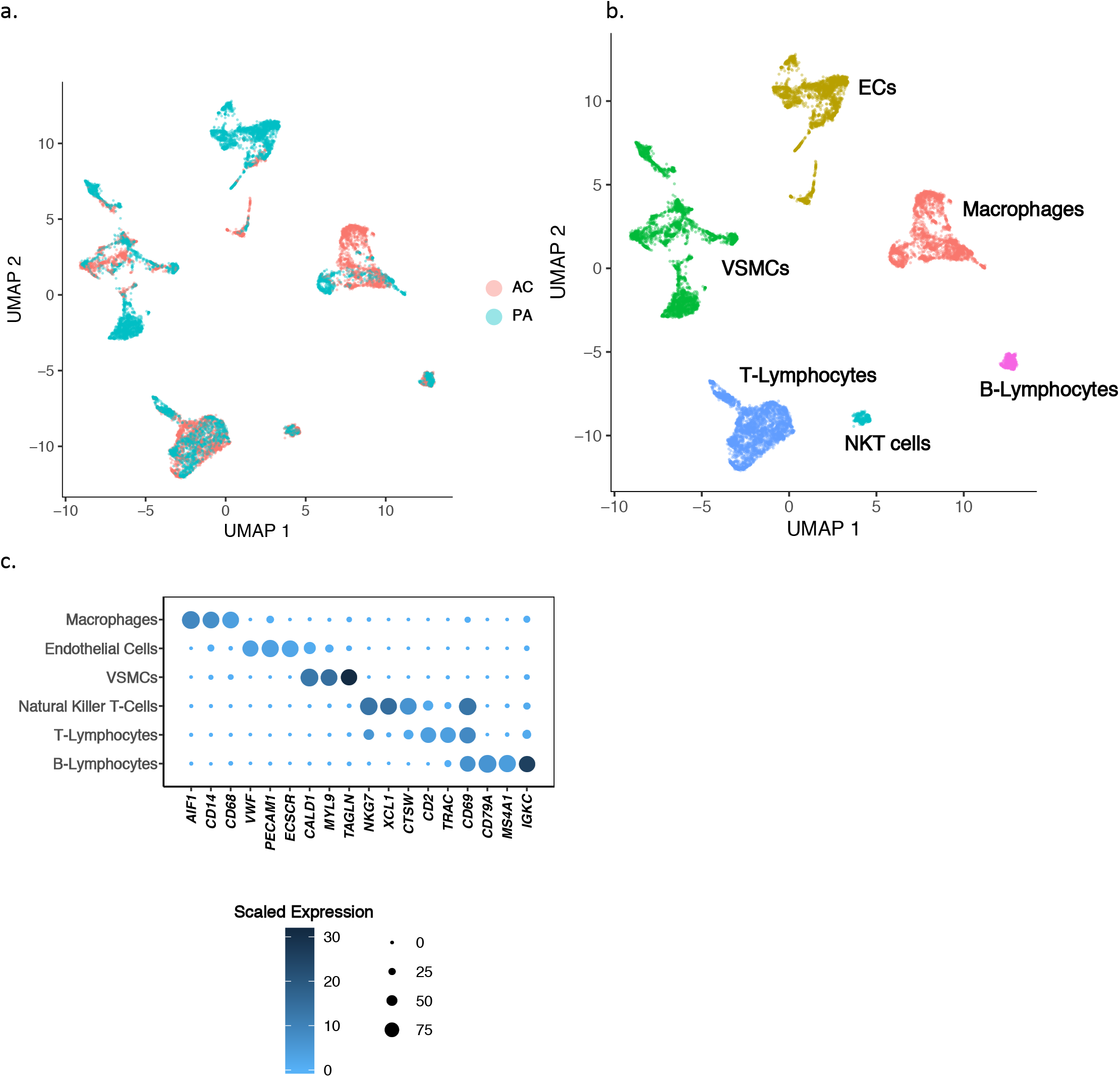
Quality control checks produce 6 main cell group clusters. **a,b** UMAP visualization of 6 major cell types following doublet removal via gene exclusion criteria (see Methods), separated by anatomic location **(a)**, and by cell type **(b)**. **c,** Dotplot depicting cell-type marker genes, resulting in the identification of macrophages, ECs, VSMCs, NKT cells, T- and B- Lymphocytes. Dot size depicts the fraction of cells expressing a gene. Dot color depicts the degree of expression of each gene.

**Fig. 3:**
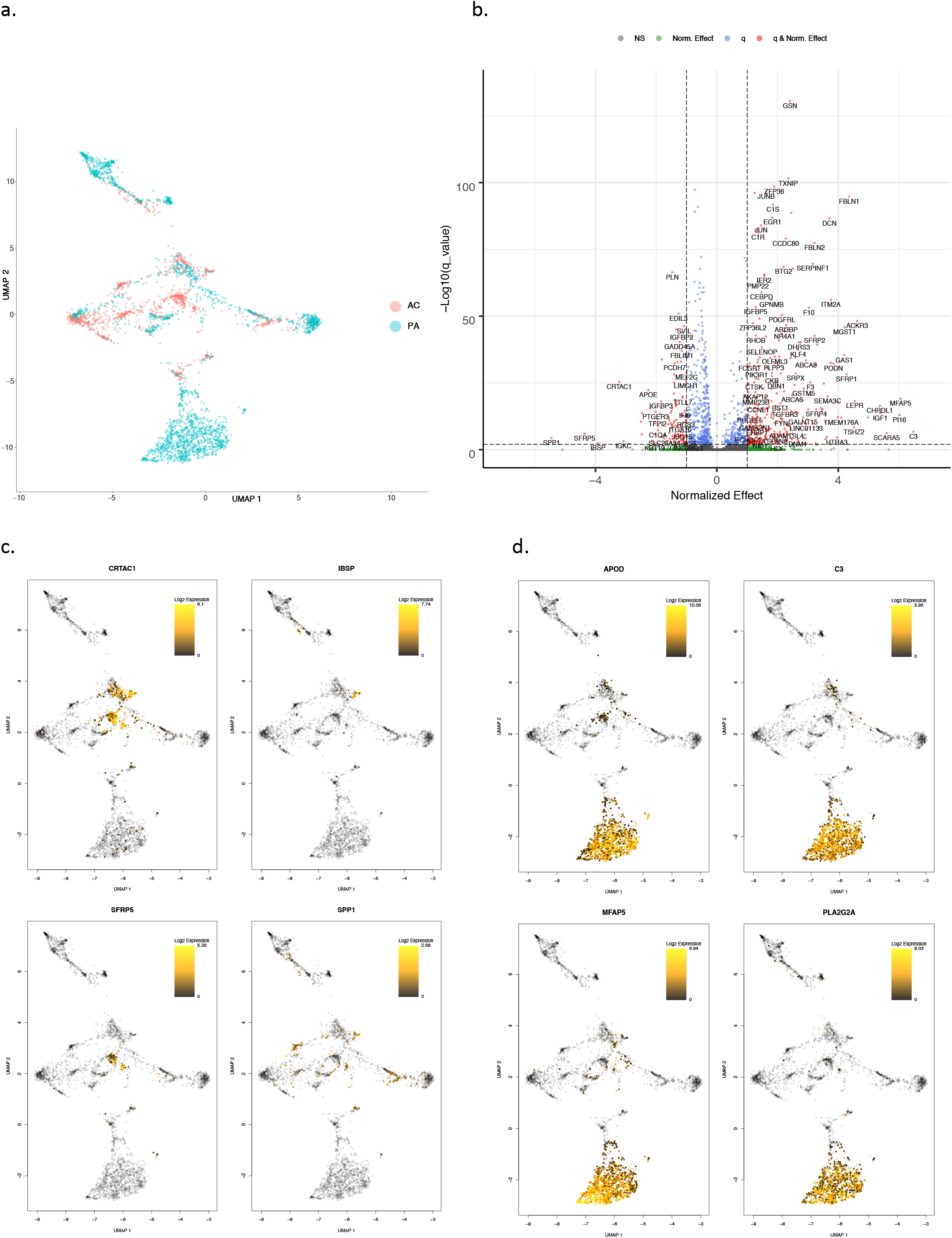
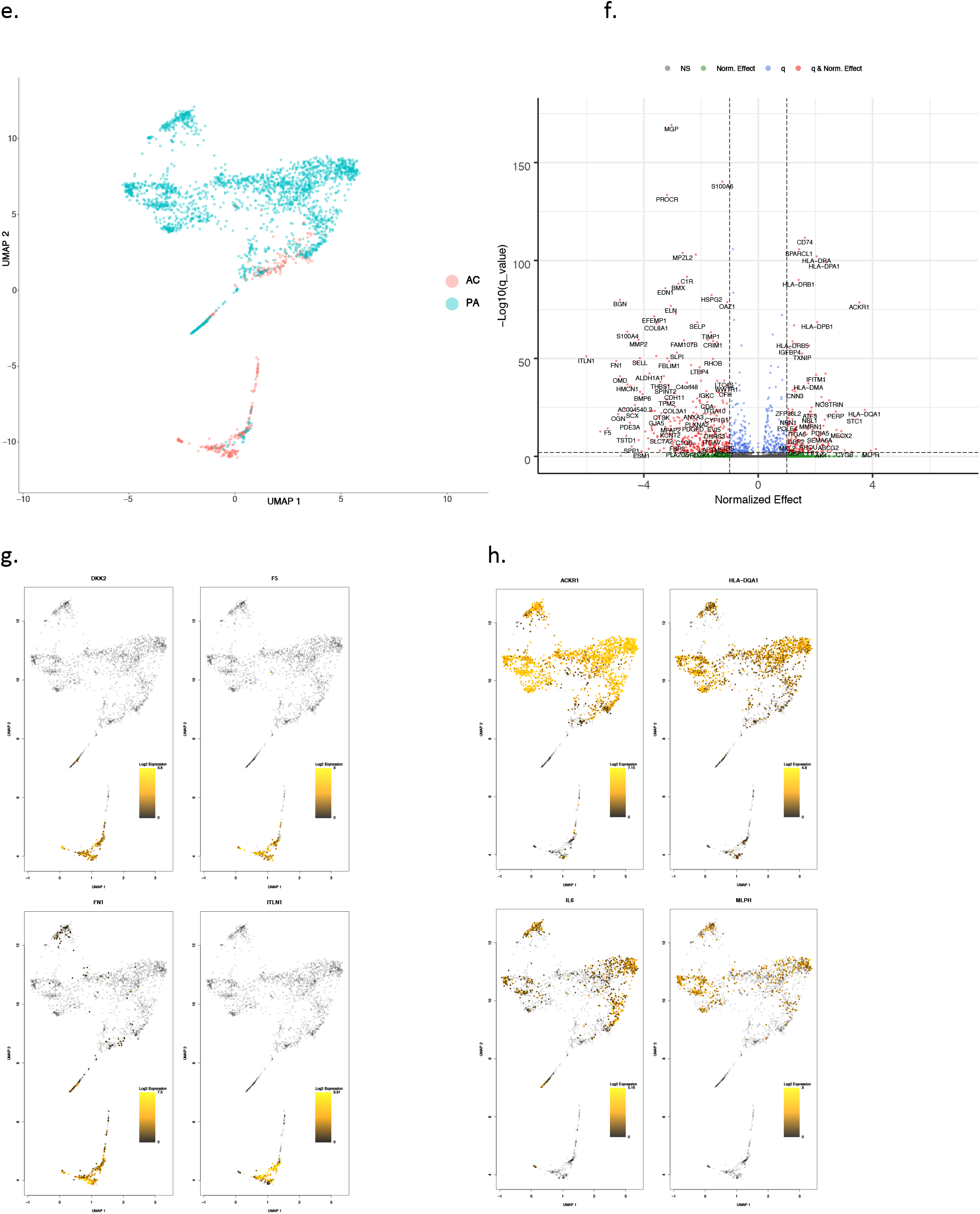
Differential gene expression in VSMCs and ECs. **a,e** UMAP visualization of VSMCs (a) and ECs (e), separated by anatomic location. **b,f** Volcano plots of the top differentially expressed genes in VSMCs (b) and ECs (f). Dotted lines represented q-value<0.01 and normalized effect >0.5 and <−0.5. **c,d** UMAP visualization of the top 4 upregulated genes in AC VSMCs (c), and PA VSMCs (d). Color bar gradient indicates lowest (black) to highest (yellow) gene expression level. **g,h** UMAP visualization of the top 4 upregulated genes in AC ECs (g), and PA ECs (h). Color scheme is similar to the above described parameters. VSMCs=3,674 cells; ECs=2,764 cells.

### Differential gene expression – Macrophages and T -Lymphocytes

Monocyte recruitment to atherosclerotic tissue and differentiation into macrophage subtypes is central to atherogenesis^14^. Our analysis indicates anatomic clustering of macrophage subpopulations, with AC cells occupying a majority of the area on the UMAP plot, overlapping somewhat with two sub-clusters of PA macrophages, suggesting at least three major macrophage sub-populations (Extended Data Fig. 4a). The top three most upregulated genes in AC macrophages relative to PA macrophages - *APOC1, FABP5* and *APOE* – all cluster together spatially in the AC-specific section of the UMAP plot and, concordant with recent findings, represent the anti-inflammatory macrophage foam cell subgroup as indicated by the high expression of *APOC1* and *APOE*^13^ (Extended Data Fig. 4c). The top three most upregulated genes in PA macrophages relative to AC macrophages include *LYVE1, SCN9A* and *S100A12* (Extended Data Fig. 4b) – forming two distinct PA macrophage sub-clusters. *LYVE1 and SCN9A* define one subset of arterial resident macrophages known to modulate arterial tone through smooth muscle collagen degradation^15,16^. The absence of these cells in the AC indicates dysfunctional arterial homeostasis in this region. *S100A12* (calgranulin c), implicated in coronary plaque formation^17^, defines a second subset of inflammatory PA macrophages^18^ (Extended Data Fig. 4c). Thus, we identify 3 macrophage subpopulations: foam cells which are the dominant population in the AC, and PA macrophages consisting mostly of inflammatory macrophages alongside a smaller group of resident PA macrophages.

In contrast to macrophages, AC and PA T-lymphocytes are spatially intermixed in the UMAP plot, indicating an overlap of cell types and states between the two anatomic regions (Extended Data Fig. 5a). Concordantly, minimal differential expression was observed between PA and AC T-cells (Extended Data Fig. 5b). The top three upregulated genes in AC-derived and PA-derived T-cells are *IGKC, MGP*, and *IGLC2,* vs *PTGDS, FGFBP2*, and *KLRF1*, respectively. *MGP, IGKC* and *IGLC2* are distributed broadly across the main body of the UMAP plot, with *MGP*, an inhibitor of ectopic tissue calcification, indicating T-cell response to calcium deposition in the AC (Extended Data Fig. 5c). Genes (*KLRF1, FGFBP2,* and *PTGDS)* upregulated in PA T-cells cluster within an offshoot off the main body of the UMAP plot suggesting a progression towards terminally differentiated effector memory T-cells in the PA. *FGFBP2* is associated with T-cell effector functions^19^, and *KLFR1* is a marker for CD4+ T-cell exhaustion and decline of cytokine releasing potential^20^. Its overexpression in the PA region suggests antigen-persistence occurs anatomically in atherogenesis. In fact, our results below demonstrate that ECs in the PA upregulate MHC class II antigen presentation genes likely resulting in this anatomic distribution of CD4+ T-cell activation and exhaustion. Full differential expression results for both macrophages and T-cells are provided in Supplementary Tables 3 and 4, respectively.

### B-Lymphocytes and NKT cells

Overall there were far fewer significant differentially expressed genes in both the B-cell and NKT populations, possibly indicating little phenotypic diversity across the two anatomic regions. Full differential expression results for both B-cells and NKT cells are provided in Supplementary Tables 5 and 6, respectively.

### Differential gene expression – VSMC and EC

GWAS results have highlighted biological processes in the vessel wall as key drivers of coronary artery disease (CAD)^21^. Further, our own work has demonstrated the vascular wall to be involved in the most impactful genetic risk factor for CAD^22^. Our results here also demonstrate extensive differential expression in these cell types across anatomic locations. Therefore, we chose to focus our efforts on dissecting expression alterations in VSMCs and ECs in order to illuminate novel pathogenic genomic signatures within these cell-types. As above, each cell type is compared across anatomic location (Fig. 3a,d), and the top differentially expressed genes are shown (Fig. 3b,e), revealing interesting spatial and expression magnitude differences between AC and PA cells.

VSMCs generate three subclusters in the UMAP plot. A large fraction of PA VSMCs reside in a subcluster occupied only by other PA VSMCs. In contrast, AC VSMCs form 2 separate clusters both which are intermingled with PA VSMCs. This suggests VSMCs occupy three major cell states, including one completely distinct from, and two that are similar to, AC VSMCs (Fig. 3a). The top four upregulated genes in the AC are sparsely expressed and include *SPP1, SFRP5, IBSP,* and *CRTAC1* (Fig. 3b,c), while *APOD, PLA2G2A, C3*, and *MFAP5* are upregulated in many PA VSMCs (Fig. 3b,d).

The spatial clustering of upregulated genes in AC VSMCs suggests the presence of separate subpopulations of matrix-secreting VSMCs involved with ECM remodeling (Fig. 3c). *SPP1* (osteopontin) is a secreted glycoprotein involved in bone remodeling^23^ and has been implicated in atherosclerosis for inhibiting vascular calcification and inflammation in the plaque milieu^24^. *IBSP* (bone sialoprotein) is a significant component of bone, cartilage and other mineralized tissues^25^. *CRTAC1* is a marker to distinguish chondrocytes from osteoblasts and other mesenchymal stem cells^26,27^. These findings suggest the presence of cartilaginous and osseous matrix-secreting VSMCs in the AC region. *SFRP5*, an adipokine that is a direct *WNT* antagonist, reduces the secretion of inflammatory factors^28^ and is thought to exert favorable effects on the development of atherosclerosis^29^. The high expression of *SFRP5* in the AC suggests a deceleration of these inflammatory processes in the core of the plaque, and an overall shift in the AC to cellular transdifferentiation, calcification and matrix remodeling.

Conversely, the upregulated genes in PA VSMCs are more ubiquitously expressed by VSMCs in a PA-specific region of the UMAP plot (Fig. 3d) and display a mixed inflammatory/anti-inflammatory profile. *C3* is highly differentially expressed in many PA cells (Fig. 3g). Complement activation has long been appreciated for its role in atherosclerosis^30^, with maturation of plaque shown to be dependent, in part, on C3 opsonization for macrophage recruitment and stimulation of antibody responses^31^. Its predominance in our PA samples suggests the role of complement activation in atherosclerosis is driven by VSMCs in the developing edge of plaque. *PLA2G2A* (phospholipase A2 group IIA), also selectively expressed by this group of cells, is proatherogenic, modulates LDL oxidation and cellular oxidative stress, and promotes inflammatory cytokine secretion^32^, further facilitating the inflammatory properties of this group of VSMCs. In contrast, *APOD* (an anti-oxidant involved with lipid uptake) decreases VSMC proliferation^33^ and has an overall cardioprotective role in CAD^34^. This combination of genes suggests a cascade of pro-inflammatory and anti-inflammatory VSMC signaling at the developing edge of plaque. Full differential expression results for VSMCs are provided (Supplementary Table 7).

Overall, we identify a gradient of increased calcification and ECM remodeling by VSMCs in the AC vs both pro-inflammatory and anti-inflammatory signaling, including recruitment of macrophages, by VSMCs in the PA.

In contrast to VSMCs, for ECs we observe a more complete separation of cells into two distinct subgroups (Fig. 3e). PA ECs significantly outnumber the AC ECs (2,316 vs 448 cells, respectively), likely due to intimal erosion during advanced disease^5^ resulting in fewer captured ECs in the AC. In addition, and in contrast to VSMCs, there is a skew towards higher magnitude expression changes in AC ECs vs PA ECs. The top four upregulated genes are *ITLN1, DKK2, F5* and *FN1* in the AC and *IL6, MLPH, HLA-DQA1*, and *ACKR1* in PA ECs (Fig. 3f).

The upregulated genes in AC ECs again suggest a synthetic profile with less overall inflammation. *ITLN* (omentin) is an adipokine enhancing insulin-sensitivity in adipocytes^35^. Interestingly, circulating plasma omentin levels were shown to negatively correlate with carotid intima-media thickness^36^, inhibit TNF-induced vascular inflammation in human ECs^37^, and promote revascularization^38^, suggesting an anti-inflammatory and intimal repair role in AC ECs. *DKK2* further indicates intimal repair as it stimulates angiogenesis in ECs^39^. The significant upregulation of *FN1* (fibronectin) in this group further suggests active ECM remodeling and may serve as a marker for mesenchymal cells and EMT-related processes^40^.

Similarly to PA VSMCs, the upregulated genes in PA ECs suggest an overall inflammatory profile. Central players in inflammation and antigen presentation are upregulated specifically in PA ECs (Fig. 3h). *IL6*, a key inflammatory cytokine associated with plaque^41^, is the most upregulated gene. Furthermore, *ACKR1,* highly upregulated in many PA ECs, binds and internalizes numerous chemokines and facilitates their presentation on the cell surface in order to boost leukocyte recruitment and augment inflammation^42^. Antigen presentation on ECs via *HLA-DQA1* (MHC class II molecule) supports activation and exhaustion of CD4+ T-cells^43,44^ as previously described. Full differential expression results for ECs are provided (Supplementary Table 8).

Overall, we identify two main EC subtypes: synthetic ECs in the AC that appear to participate in intimal repair, revascularization, and ECM modulation, and inflammatory ECs in the PA region that facilitate inflammation via antigen/chemokine presentation and recruitment of immune cells, including CD4+ T-cells.

### Hallmark Processes – VSMCs and ECs

In order to explore the anatomic differences for these cell types further, gene set enrichment analysis (GSEA) was used to asses hallmark processes most significantly altered in VSMCs and ECs (Fig. 4a,b). Epithelial to Mesenchymal Transition (EMT), oxidative phosphorylation, and myogenesis gene upregulation were strongly enriched in both AC VSMCs and ECs, collectively suggesting an increase in cellular metabolic activity and proliferation as they participate in transdifferentiation. In contrast, a distinctly inflammatory profile was seen in PA VSMCs and ECs, with IFN gamma/alpha responses and TNFa signaling via NFkB dominating the enriched processes in these groups of cells. Because EMT and TNFa signaling were both shared and strongly enriched processes in the two cell types, the gene signatures associated with these hallmarks were further scrutinized through generation of heatmaps consisting of leading-edge differentially expressed genes from each hallmark process (EMT - Fig. 4c, TNFa signaling via NFkB - Extended Data Fig. 6a).

**Fig. 4:**
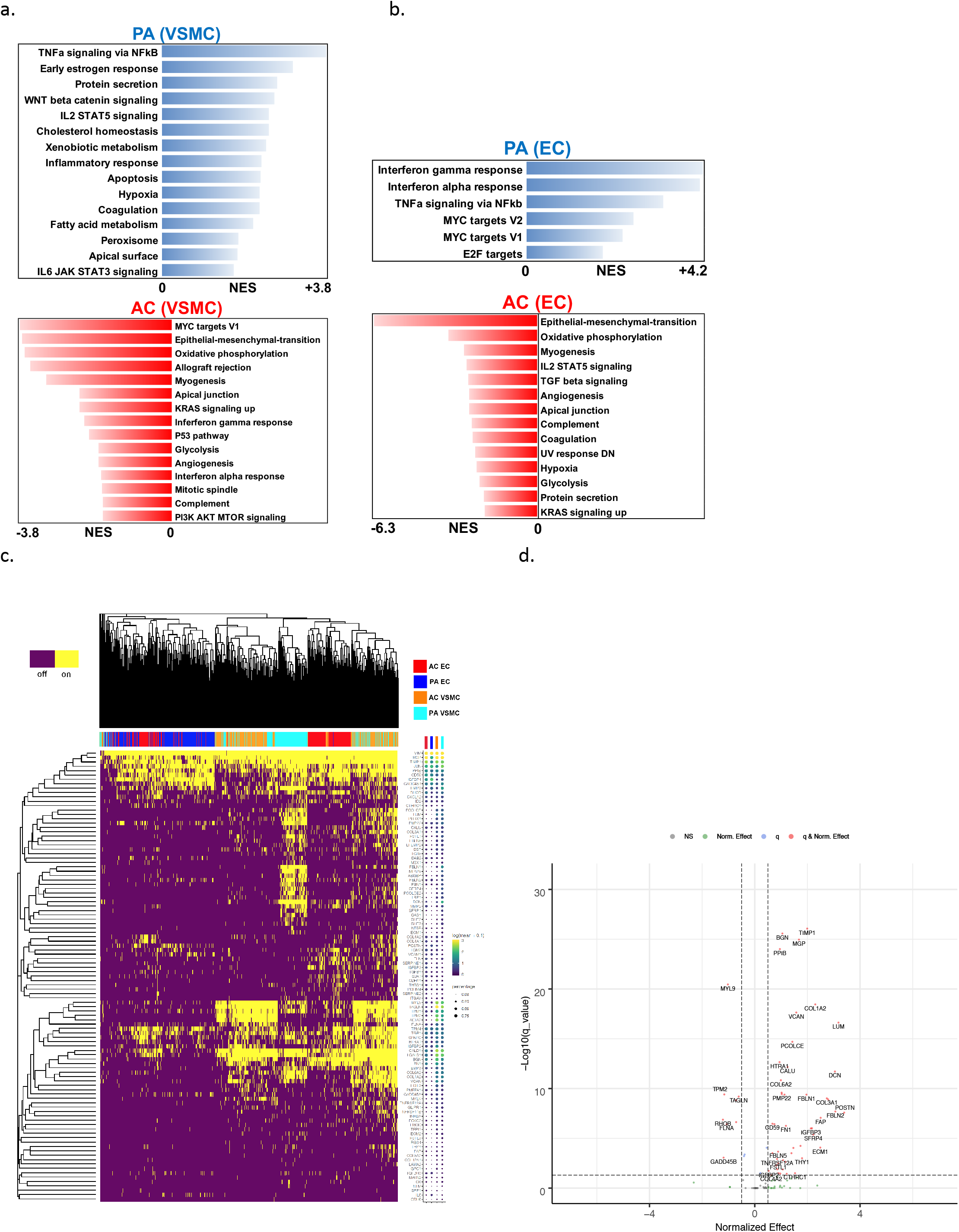
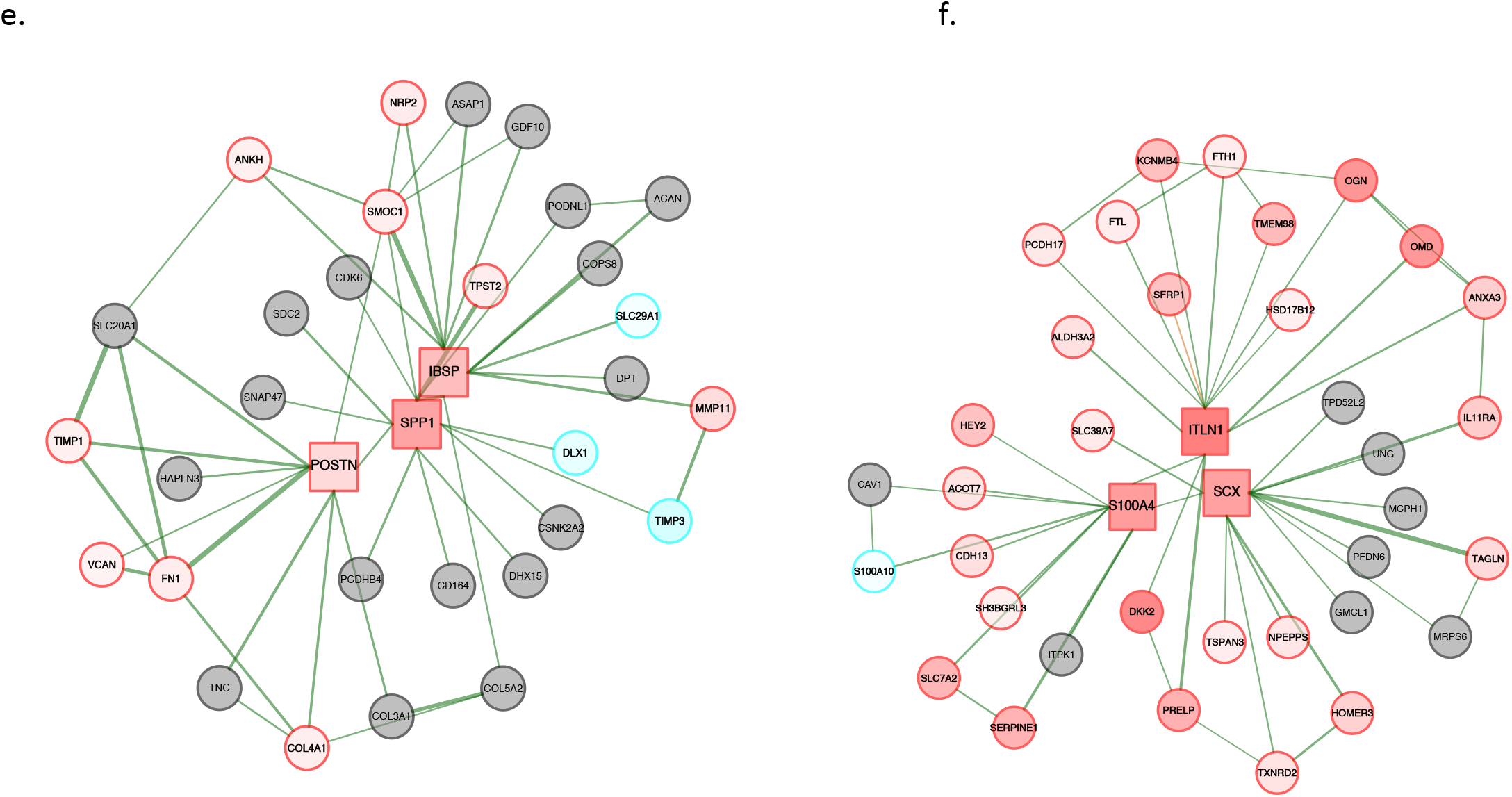
Gene set enrichment analysis and gene co-expression networks identify key gene drivers of EMT hallmark biologic process. **a,b** Normalized enrichment score (NES) ranking of all significant PA and AC Hallmarks generated from GSEA analysis of differentially expressed genes for VSMCs (a) and ECs (b) (FDR q-value<0.05). **c,** Fully clustered on/off heatmap visualization of overlap between leading edge EMT hallmark genes generated by GSEA. Heatmaps are downsampled and represent 448 cells from each cell type and anatomic location (1792 total cells). A dotplot corresponding to gene expression levels for each cell type in the heatmap is included. Dot size depicts the fraction of cells expressing a gene. Dot color depicts the degree of expression of each gene. **d,** Volcano plot of differentially expressed genes between the two groups of dysfunctional VSMCs in (c). Dotted lines represented q-value<0.01 and normalized effect >0.5 and <−0.5. **e,f** Gene co-expression networks generated from VSMC Module 13 (d) and EC Module 1 (e) representing the EMT hallmark from GSEA analysis. Genes are separated by anatomic location (red=AC genes, cyan=PA genes), differential expression (darker shade=higher DE, gray=non-significantly DEGs), correlation with other connected genes (green line=positive correlation, orange line=negative correlation) and strength of correlation (connecting line thickness). Significantly DEGs (q<0.05) with high connectivity scores (>0.3) are denoted by a box instead of a circle.

**Fig. 5:**
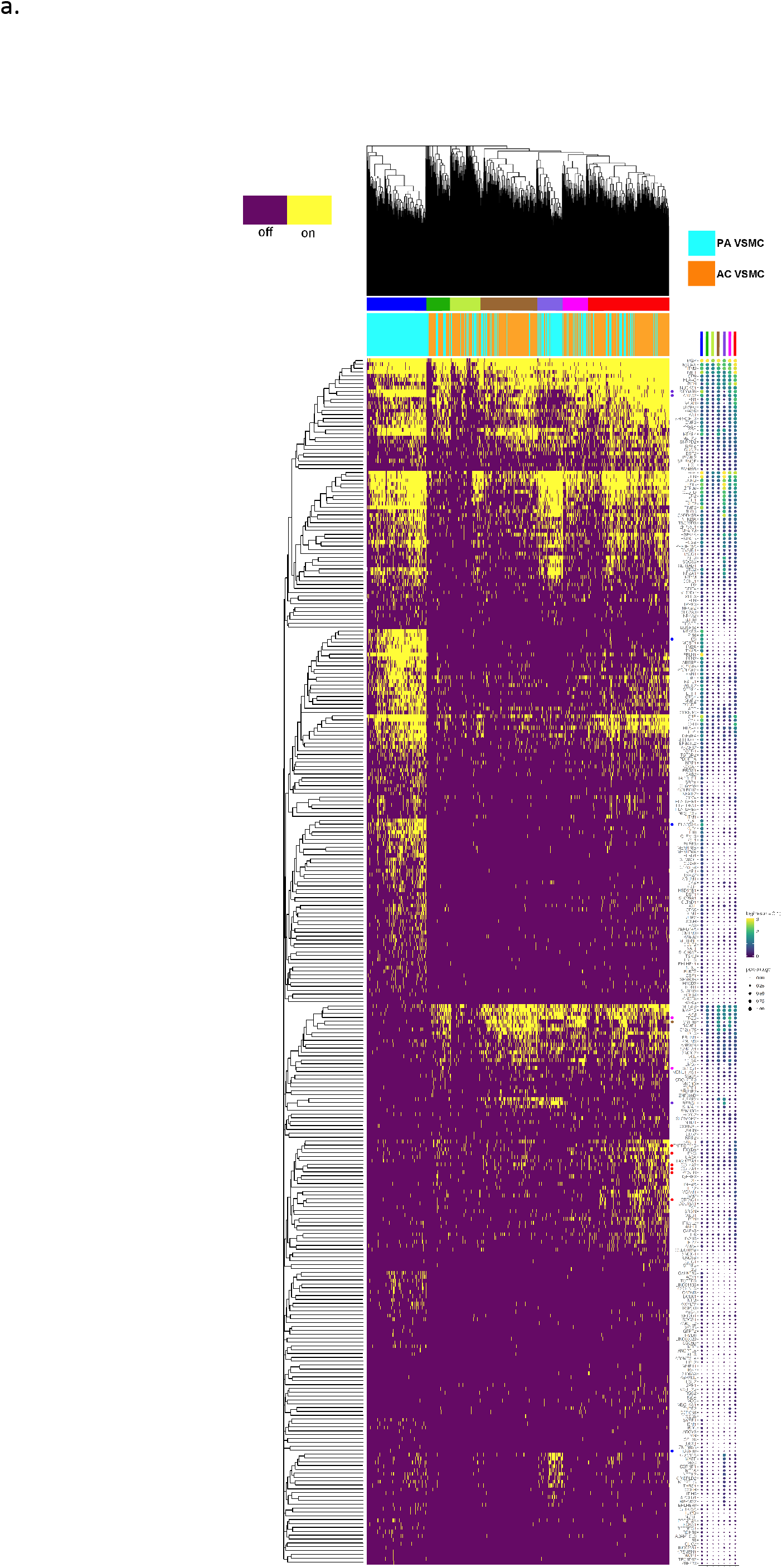
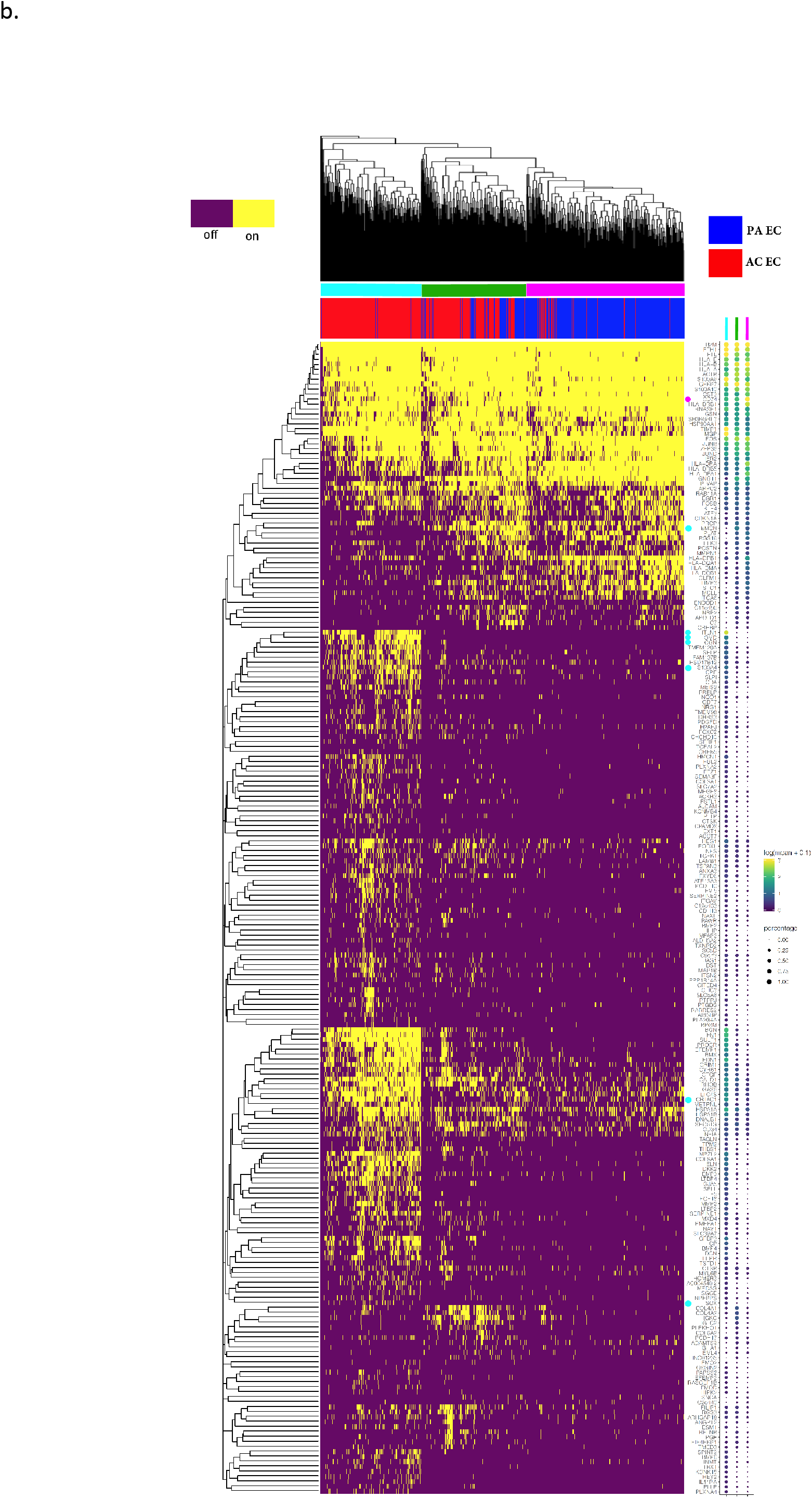
Evaluation of VSMC and EC subpopulations. **a,b** Biclustered heatmap visualization of all significant genes (q<0.05) from VSMC (a) and EC (b) modules enriched with differentially expressed genes. (a) 1224 VSMCs from each anatomic location (2448 cells total). Large color bar denotes PA (cyan) and AC (orange) VSMCs. Small color bar above denotes distinct cell subpopulations (blue, forest green, lime green, brown, purple, magenta, red). (b) 448 ECs from each anatomic location (896 cells total) in. Large color bar denotes PA (blue) and AC (red) ECs. Small color bar above denotes distinct cell subpopulations (cyan, green, magenta). A dotplot corresponding to gene expression levels for each cell subpopulation on the heatmap is included. Colored dots next to specific genes correspond to critical genes related to the designated cell subpopulation.

**Fig. 6:**
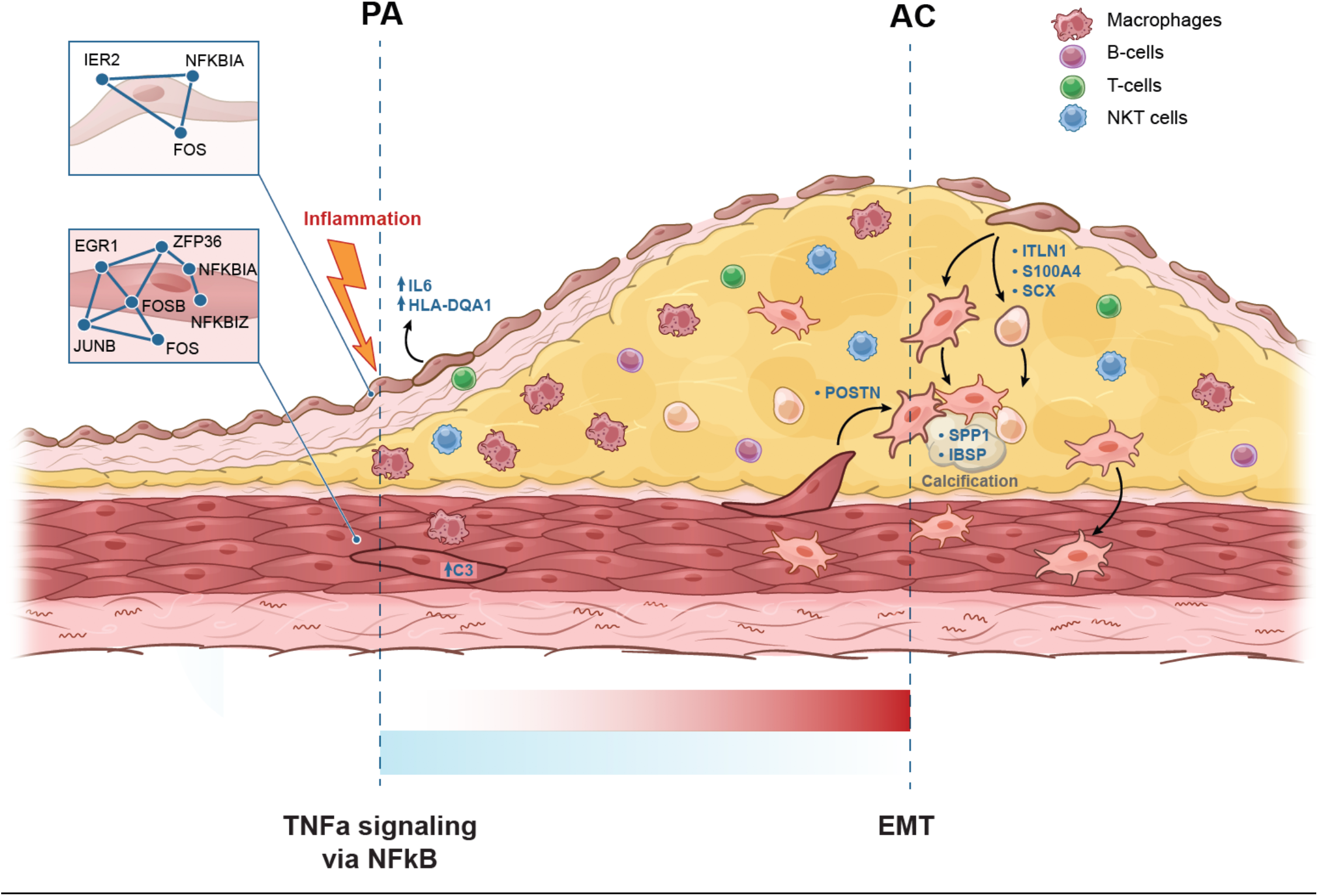
Schematic illustrating key hallmark processes in PA and AC VSMCs and ECs. See main text for details.

While overlapping at the hallmark level, separation of cells by cell type as well as anatomic location in the EMT hallmark heatmap suggests the overlapping processes are mediated by distinct gene sets in each cell type. Moreover, analysis of EMT hallmark genes further supports the presence of 2 cellular subtypes of AC VSMCs as they appear to cluster into two distinct groups of cells with dichotomous expression of contractile *(MYL9, TPM2, TAGLN, FLNA*) versus synthetic/CTD (*POSTN, LUM, FBLN2, DCN, PCOLCE2, MGP, COL3A1*) gene signatures (Fig. 4c,d). VSMC plasticity is a long appreciated phenomenon in vascular pathophysiology^45^, and these results indicate a group of VSMCs in the AC perform the contractile functions of the blood vessel wall, while the other group of VSMCs perform CTD, proliferate, migrate, and remodel the ECM.

In contrast to distinct subclustering of cells by CTD genes, there appears to be a common gradient of genes involved in inflammation and response to inflammation expressed throughout the atherosclerotic tissue, with higher levels of TNF-related inflammatory genes expressed in PA VSMCs and ECs compared to AC cells, indicating a predominance of inflammatory processes occurring in the developing edge of plaque overall (Extended Data Fig. 6a). Collectively these genes (*EIF1, FOS, JUN, JUNB, ZFP36, PNRC1, KLF2, IER2, CEBPD, NFKBIA, GADD45B, EGR1, PPP1R15A* and *SOCS3*), in addition to IL6-driven signaling in PA ECs (Fig. 3f), appear to coordinate the inflammatory response pathways in plaque and its adjacent structures. All cell types analyzed thus far are coordinated along this gradient of inflammation.

### Network Analysis

To identify the key genes driving the differentially expressed hallmark processes observed in ECs and VSMCs we reconstructed gene co-expression networks using a novel partial correlation-based approach (see *Methods*), defined modules by clustering, and overlaid differential expression analysis results on these modules to identify those enriched in genes differentially expressed between AC and PA tissues.

Using this strategy, 31 and 39 distinct gene network modules were generated in our VSMC and EC datasets, respectively (Extended Data Fig. 7,8). Of these, 7 modules in VSMCs (Modules 9, 11, 13, 24, 28, 30 and 31), and 5 modules in ECs (Modules 1, 2, 6, 35, and 36) were enriched with differentially expressed genes (p-value<0.05, hypergeometric test, see *Methods*). Differentially expressed EMT/CTD hallmark genes overlapped significantly with module 13 of VSMCs and module 1 of ECs. Differentially expressed TNFa signaling via NFkB hallmark genes overlapped significantly with module 31 of VSMCs and 36 of ECs (p-value<0.05, hypergeometric test).

### CTD in VSMCs and ECs from the Atherosclerotic Core

In order to further characterize genes critical to stimulating CTD in AC VSMCs and ECs we examined gene co-expression networks in conjunction with differential expression data from the modules enriched with EMT hallmark genes. In VSMC module 13 we identified 9 genes (*SPP1, IBSP, POSTN, MMP11, COL15A1, FN1, COL4A1, SMOC1, TIMP1*) whose expression was significantly upregulated in AC cells and with strong network connectivity (see *Methods*). Among these genes we identify *POSTN*, *SPP1* and *IBSP* as key gene drivers of CTD in AC VSMCs due to their strong central connectivity and high degree of differential expression in the CTD network (module 13) (Fig. 4e). *POSTN* (periostin) is expressed by osteoblasts and other connective tissue cell types involved with ECM maturation^46^ and stabilization during EMT in non-cardiac lineages^47,48^. *POSTN, SPP1,* and *IBSP* are highly interconnected in our network and likely serve as drivers of CTD by modulating other correlated genes such as *TIMP1, VCAN, TPST2, SMOC1, MMP11, FN1,* and *COL4A1* (Fig. 4e), all genes which are involved with cellular differentiation^49^ and extracellular matrix remodeling^50,51^.

In module 1 of our EC network we identified 18 genes (*ITLN1, FN1, OMD, S100A4, SCX, PRELP, GDF7, TMP2, SERPINE2, SLPI, HEY2, IGFBP3, FOXC2, RARRES2, PTGDS, TAGLN, LINC01235, and COL6A2*) whose expression was significantly upregulated in AC cells and with strong network connectivity. Among these genes we identify *ITLN1, S100A4* and *SCX* as key gene drivers of CTD in ECs associated with the AC (Fig. 4f). *ITLN1* (omentin) is highly upregulated in ECs associated with the atherosclerotic core, and network data indicate it is strongly correlated with genes involved with cellular proliferation and ECM modulation. *ITLN* is also strongly correlated to *OGN* (osteoglycin) which induces ectopic bone formation^52^, indicating that *ITLN1* modulates ECs with osteoblast-like features in the atherosclerotic core. *SCX* (scleraxin), a transcription factor that plays a critical role in mesoderm formation and the development of chondrocyte lineages^53^, is co-expressed with *IL11RA*, an interleukin receptor implicated in chondrogenesis^54^, as well as with a variety of genes involved with cellular development and modulation of ECM structures. Thus, *SCX* modulates chondrocyte-like ECs in the AC. *S100A4* is a calcium-binding protein that is highly expressed in smooth muscle cells of human coronary arteries during intimal thickening^55^, and silencing this gene in endothelial cells prevents endothelial tube formation and tumor angiogenesis in mice^56^. Co-expression with *HEY2*, a transcription factor involved with *NOTCH* signaling and critical for vascular development^57^, indicates an important role in repair via re-endothelialization of plaque-denuded artery.

### TNFa Signaling via NFKB in Proximally Adjacent VSMCs and ECs

Next, genes critical to stimulating TNFa signaling via NFkB in PA VSMCs and ECs were evaluated. In VSMC module 31 we identified 14 genes (*APOLD1, MT1A, ZFP36, EGR1, JUNB, FOSB, JUN, FOS, RERGL, MT1M, DNAJB1, CCNH, HSPA1B* and *HSPA1A*) whose expression was significantly upregulated in PA cells and with strong network connectivity. Among these genes we identify immediate-early (IE) genes *ZFP36, EGR1, JUNB, FOSB* and *FOS* as critical response genes in this hallmark process which cluster tightly together and correlate with *NFKBIA* and *TNFAIP2* - both inhibitors of NFkB-mediated signaling^58^ (Extended Data Fig. 6c). The presence of these genes suggests an IE-mediated anti-inflammatory response to inflammation in the PA region, which is eventually lost leading to CTD.

In EC module 36 we identified two genes (*IER2* and *FOS*) whose expression was significantly increased in PA EC cells (Extended Data Fig. 6e), and are highly correlated with other critical transcription factors in our network, including *FOSB, JUNB, EGR1* and *ZFP36*, further supporting this group of gene’s importance in the TNFa signaling hallmark (Extended Data Fig. 6e). Similar to VSMCs, these genes are also correlated with *NFKBIA* in the network, again suggesting an IE-mediated anti-inflammatory response to inflammation in the PA region, eventually leading to CTD.

### Evaluation of Cellular Subpopulations

Finally, in order to identify and characterize refined subpopulations from each anatomic region, we selected the 7 VSMC and 5 EC differentially expressed modules described above and biclustered cells and genes (Fig. 5a,b). Groups of cells were then correlated with gene networks in order to identify biologically distinct cell subpopulations.

### VSMCs

We identified seven distinct populations of VSMCs with some overlapping features in our analysis (Fig. 5a). The first subpopulation (Fig. 5a, red bar) has elevated expression of *POSTN* (osteoblasts; NE=-2.206, q= 3.60e-16)*, CRTAC1* (chrondrocytes; NE=−3.22, q= 3.91e-26), *TNFRSF11B* (bone remodeling; NE=−0.98, q= 7.31e-06)^59^, *ENG* (VSMC migration; NE=0.87, q= 1.41e-13)^60^, *COL4A2* and *COL4A1* (cell proliferation, association with CAD; NE=−0.98, −1.03 and q= 3.17e-15, 5.68e-11, respectively)^61,62^, and a variety of interferon alpha inducible (IFI) genes which modulate numerous cellular functions from proliferation to apoptosis. *SFRP5* (anti-inflammatory; NE=−4.36, q=9.84e-07) is upregulated almost exclusively in this population of VSMCs. Collectively this group of cells represents terminally transdifferentiated osteoblast- and chondrocyte-like VSMCs which facilitate calcification and cartilaginous matrix-secretion and reside largely in the AC.

The adjacent VSMCs group (Fig. 5a, magenta bar) is similar to the above described population, but consists of a larger fraction of PA VSMCs, indicating that these cells are in the process of completing the transition to osteoblast- and chondrocyte-like VSMCs. Key gene differences between these two groups includes comparatively higher levels of *FRZB* (may regulate chondrogenesis; NE=−0.54, q=1.93e-07)^63^, *SCRG1* (stimulator of chondrogenesis; NE=−0.73, q=0.032)^64^, and lower levels of *CRTAC1* (chondrocyte marker)^27^, again indicating these cells are in the process of transitioning to a chondrocyte-like phenotype. *SCRG1* is strongly co-expressed with *CYTL1* (regulation of chondrogenesis)^65^, further cementing this group of cells as chrondrocyte-like VSMCs in the AC.

The next cluster on our heatmap consists of mostly PA VSMCs (Fig. 5a, purple bar). These are likely PA VSMCs in the early stages of CTD and eventually arriving at the AC-state given their overall similarity in expression to the above two subpopulations. These cells display a robust inflammatory IE response. Interestingly, this group displayed the highest level of *RERGL* expression (NE=1.15, q=0.00012), a member of the RAS family of GTPases that has been liked to lymphovascular invasion in breast cancer^66^. Thus, these cells are likely responding to inflammation, and are likely in the process of migration and CTD, falling in between inflammatory and terminally transdifferentiated cells on our continuum.

The next population (Fig. 5a, brown bar) is comprised of contractile AC VSMCs expressing *MYH10* (NE=−1.0, q= 4.20e-27) which, as described earlier, support the contractile functions of the blood vessel wall.

The next two populations (Fig. 5a, forest and lime green bars) represent a mixture of VSMCs with overall pan-decreased expression levels compared to other subpopulations. It is unclear what unique function, if any, these groups of cells possess.

The final identified population consists exclusively of PA VSMCs (Fig. 5a, blue bar). This group expresses higher levels of IE genes relative to all AC VSMCs and represents cells involved in and responding to inflammation, immune cell recruitment, and the initiation of CTD. *C3* (opsonization and macrophage recruitment; NE=6.5, q=1.74e-07) is highly differentially expressed in this cohort and thereby augments PA atherogenesis through inflammation and macrophage recruitment. *PLA2G2A* (promotes inflammatory cytokine secretion; NE=7.25, q=0.046) is also selectively expressed by this group of cells, further facilitating their inflammatory properties. This group of VSMCs also shows early migratory and CTD-like qualities given the expression of *FBN1, SEMA3C, HTRA3* and *C1QTNF3*, (NE=2.77, 3.65, 4.0, 3.58, respectively; q=6.93e-41, 1.25e-20, 2.53e-05, 0.00012, respectively) genes that are both highly differentially expressed in this cohort and with high signal strength in our networks (Fig. 5a, Extended Data Fig. 7a). *FBN1* (ECM component) is strongly correlated with *TGFBR3, SEMA3C* and *CD248* (modulators of EMT-like processes)^67–69^. Interestingly, this group of cells co-expresses *IGSF10*, a marker of early osteochondroprogenitor cells^70^, *TMEM119* (bone formation and mineralization; promotes differentiation of myeloblasts into osteoblasts)^71,72^ and *WNT11* (bone formation)^73^ (Extended Data Fig. 7a). Collectively this population of PA VSMCs represents cells responding to and promoting inflammation at the developing edge of the plaque and initiating the early stages of CTD.

Overall, there are three major VSMC subpopulations, each of which show transitory properties resulting in subpopulations transitioning through the inflammatory to CTD continuum. First, there are two groups of inflammatory PA VSMCs that are involved in recruitment of inflammatory mediators and show early signs of CTD. Second, a group of contractile mostly AC VSMCs that support normal functions of the vascular wall. Finally, two groups of also mostly AC synthetic osteoblast- and chondrocyte-like VSMCs at various stages of transdifferentiation.

### ECs

We identified two distinct populations of ECs and one with overlapping features with the other two in our analysis (Fig. 5b). The majority of ECs from the AC form one large cluster of osteoblast- and chondrocyte-like ECs (Fig. 5b, cyan bar) largely devoid of endothelial-marker gene *EMCN*^74^ (NE=0.86, q=1.17e-09). Critical EMT genes identified earlier (*ITLN1*, *SCX* and *S100A4*) are predominantly expressed in this large cluster of AC ECs alongside highly correlated genes *OMD, OGN*, and *CRTAC1* (Fig. 5b, 4f), again indicating that this population of ECs represents the main group of transdifferentiated ECs.

The next group of cells represents a mixture of AC and PA ECs with similar gene signatures at varying levels to each of the other two groups of cells (Fig. 5b, green bar), likely representing dysfunctional ECs that are in transition from the below described inflamed subtype to the above described CTD subtypes.

The final group of cells represents the inflammatory PA subset of ECs (Fig. 5b, magenta bar). This group has a greater number of cells expressing immune genes such as the cluster of HLA genes, as well as *CD74* (NE=1.63, q=2.07e-112), a gene which forms part of the invariant chain of the MHC II complex and is a receptor for the cytokine macrophage migration inhibitory factor (MIF)^75^. The upregulation of MHC class II complex in this subset of PA ECs complements our previous finding of CD4+ T-cell recruitment to this subpopulation of PA ECs, leading to over-activation and exhaustion via antigen-persistence.

Overall, there are two main EC subpopulations, and similar to VSMCs these cells display transitory properties as they move through the continuum, resulting in three identified cell states. First, there is a group comprised near exclusively of inflammatory PA ECs that is involved in recruitment of inflammatory mediators. Second, we identify a near exclusive group of synthetic osteoblast- and chondrocyte-like AC ECs. Lastly, there is an intermixed group of ECs with varying degrees of gene expression related to both inflammation and CTD on the identified continuum.

## Discussion

In this study, unbiased single-cell transcriptomics of type VII calcified plaque revealed 15 distinct cell types, including six major cell types; macrophages, ECs, VSMCs, NKT cells, T- and B-lymphocytes. When expression signatures were contrasted across AC and PA plaque regions, macrophages, VSMCs, and ECs demonstrated the greatest degree of anatomical diversity. Our findings confirm the identification of three primary macrophage cell types in human plaque tissue: inflammatory, resident, and foam cells, with foam cells dominating the area of carotid artery most burdened with disease (the AC) and resident and inflammatory cells strongly enriched in the developing edge of the plaque (the PA). For VSMCs and ECs, we determine that both cell types contribute to increased calcification and ECM remodeling activity through CTD in the AC. In the PA, both pro-inflammatory and anti-inflammatory signaling including recruitment of immune cells is enriched in both cell types. However, differing biological processes and CTD molecular networks underly this convergent biology.

VSMC plasticity in response to injury and environmental cues is central to vascular homeostasis and remodeling^76^. *KLF4*-dependent modulation of phenotype switching in VSMCs was shown to be pathogenic, and deletion of this gene in a murine model of atherosclerosis reduced lesion size^77^, suggesting that targeting modulators of CTD in atherosclerosis may be an effective treatment strategy. It is therefore critical to unmask novel modulators of CTD, and to explore this phenomenon in other cells of the vascular wall such as ECs. Similarly, endothelial-mesenchymal-transition (EndMT) is an appreciated phenomenon representing a link between inflammatory stress and endothelial dysfunction^78^, but little is known about its genetic modulation, and/or whether the modulation of CTD differs in VSMCs and ECs. In our study, CTD in the AC to osteoblast- and chondrocyte-like cell states is defined by different molecular networks with different key driver genes: *POSTN, SPP1 and IBSP* in VSMCs, vs *ITLN1, SCX* and *S100A4* in ECs. While these groups of genes both promote CTD in their respective cell types, their pathogenic versus protective role are unclear. The VSMC genes generally promote ossification and may have a more deleterious role in atherogenesis than the cohort of genes promoting CTD in ECs, some of which are known to promote re-endothelialization and intimal repair. Further scrutinization of these genes will be critical to fully appreciate their contribution to disease progression.

Similarly, while inflammatory signaling is enriched in the PA for both cell types, the molecular networks engaged in inflammatory signaling differs across VSMCs and ECs. Both cell types appear to respond to inflammatory signaling via IE-mediated expression of NFkB inhibitors; *NFKBIA* and *TNFAIP2*. However, VSMCs engage in immune cell recruitment through expression of *C3* (complement component 3), whereas ECs engage in immune cell recruitment through MHC class II presentation. A wide variety of other inflammatory processes are activated in both cell types, converging on TNFa signaling via NFkB in both cell types and IL6 signaling in ECs – both likely augmented by vascular wall driven recruitment of inflammatory mediators of early plaque development^79^.

Illuminating the anatomic landscape of atherosclerosis at single-cell resolution underscores the intriguing possibility of employing site-specific therapies to curtail disease progression. Given the recent focus on targeting inflammation to impede atherogenesis^80^, our results highlight the importance of targeting inflammatory mediators at the nidus of disease, thereby halting the development of plaque and progression of blood vessel cells into pathogenic CTD. Similar strategies may be contemplated for the CTD observed in our patients, though it is unclear whether AC calcification is pathogenic or stabilizing and protective.

Overall our study provides unprecedented anatomic insight into atherogenesis by examining key cellular processes occurring in the atherosclerotic core and adjacent arterial tissue and lays the groundwork for continuing efforts to unravel the complexity of this disease.

## Methods

### Selection of patients

Identification and consenting of three patients with severe carotid plaque formation requiring carotid endarterectomy occurred in collaboration with the Scripps Health Biorepository under IRB# 19-7332 approved by the Scripps Institutional Review Board. Patient characteristics and comorbidities are presented in Supplementary Table 1. Plaques were characterized by histopathology according to AHA classification scheme^81^. Briefly, near full thickness sections of artery and plaque were recovered from the atherosclerotic core (except adventitia), and full thickness proximally adjacent arterial sections were recovered from the same patient during the cut-down portion of the endarterectomy. These specimens were immediately transported for tissue processing.

### Tissue processing and generation of single cell suspensions

AC and PA carotid artery from patients undergoing carotid endarterectomy was immediately collected from the operating room and transferred on ice for tissue dissociation (Fig 1a,b). Samples were weighed, minced with surgical blades, then placed in conical tubes with pronase (5000U/ml) and collagenase II (0.1% solution). Tissue was then incubated at 37C for one hour with continuous gentle agitation. 30min into the incubation period, DNase I solution was added, and the entire solution remained incubated for an additional 30min. Pipetting mixture up and down every 10min helped break tissue apart. After one-hour incubation was complete, 10% FBS+Complete EC media (Cell Applications, Inc) was immediately added to dissociated tissue to quench enzymatic activity. The sample was then inverted a few times and then set down for 30sec, allowing all debris to settle at the bottom. The entire solution was carefully transferred, without settled debris, to another conical tube (use pipettor to transfer solution near debris pellet carefully, removing all solution but leaving pellet at bottom of tube). Sample was then centrifuged at 500g for 5min and supernatant discarded. Cell pellet was resuspended in 5mL 1X RBC lysis buffer and incubated at RT for 5min. 30mL 1X PBS was added to stop the reaction and the sample was then spun immediately at 500g for 5min at RT. The supernatant was discarded, and cell pellet resuspended in 10mL FACS buffer composed of: 1X PBS, 2.5mM EDTA, 25mM HEPES, 1% FBS (heat-inactivated), and 1% Pen-Strep. To wash excess reagents, the sample is centrifuged again at 500g for 5min, supernatant discarded, then cell pellet is again resuspended in 2mL FACS buffer. 2ul of 5mM DRAQ5 is added to each tube, and solution is pipetted up and down a few times. Next, solution is triturated with P1000 and entire sample sieved through 100uM cell strainer fitted onto 50mL conical tube. This step is repeated next through a 40uM cell strainer. The solution is placed on ice and immediately transported for FACS. Cells were sorted on a MoFLo Astrios EQ cell sorter (Beckman Coulter) with a 100 μm nozzle tip at a sheath pressure of 20 psi. Cells were distinguished from cellular debris by gating DRAQ5 positive events and doublets excluded using appropriate FSC and SSC gating. (Extended Data Fig. 1a-d). Cells isolated represented <1% of the total particles in the sample which was then transported for evaluation of viability and subsequent sequencing.

Cells were then processed on the 10X Genomics Chromium single cell 3’ v3 gene expression platform. Cells were counted and checked for viability using a Countess II Automated Cell Counter (Thermo Fisher). Sequencing libraries were prepared as recommended in the user manual with cell target numbers ranging between 6,000 – 22,000 cells per sample. Completed sequencing libraries were then sequenced on an Illumina NextSeq500 using 28 × 91 paired-end reads with 8 base i7 index reads to demultiplex different samples. Aggregating our 6 samples (3 patient-matched PA and AC samples) resulted in 51,981 cells, a sequencing depth of 15,549 reads/cell, 1,339 median genes/cell and 3,776 UMI/cell.

### Single-Cell Transcriptomics

We used a combination of publicly available tools and custom scripts to process single-cell transcriptomic data. These methods have been developed and refined by our lab for the analysis of single-cell transcriptomics of human atherosclerotic plaque tissue, an extremely challenging tissue for single-cell sequencing given the variable degree of cellular viability and cellular and extracellular matrix heterogeneity present in plaque tissue from vascular wall to the fatty plaque tissue itself. A combination of 10X Cell Ranger^82^, Monocle^83^, Seurat^84^, and custom R and python scripts as well as pathway analysis tools are combined in a comprehensive single-cell quality control and analysis pipeline.

### Cell Set Preparations, Aggregation, and Initial Cell-Type Identification

Raw single cell sequencing data was processed independently per sample using the 10X Genomic Chromium platform^82^. 10X Genomic’s Cell Ranger with default parameters for read mapping^85^, sample quality control, unique molecular identifier filtration, normalization and expression quantification. Further quality filtering is performed by Seurat^84^ to remove cells with greater than 10% mitochondrial mRNA (total mRNA), in addition to cells with less than 200 or greater than 4,000 genes expressed. Quality control resulted in a reduction of 6,145 cells, down to 45,836 cells total prior to downsampling. Cell sets were then down-sampled to 17,100 cells to account for imbalances in total cell counts across samples which may adversely influence clustering analysis.

Aggregated cell sets were then further processed by Monocle for cell-type sub-setting^83^. Dimensionality reduction was performed as standard via principal component analysis in Monocle and UMAP partitioning is applied with the default Louvain algorithm^86^, which we have found to perform better than t-SNE. In practice, we will apply and compare both clustering approaches. Cell type assignment was a largely manual process facilitated by partition level differential gene expression analysis to identify 3 known marker genes per cell type that were expressed in greater than 80% of cells and at a mean expression count greater than 2. At this point partitions were assigned to cell types.

### Doublet analysis and filtering

In order to identify and remove distressed cells, or rare cell types and artifacts, we first estimated doublet-rates and attempted to filter doublets using the Scrublet package^87^. However, we found that Scrublet and other doublet detection packages were not effective at identifying doublets in highly heterogenous samples like atherosclerotic plaque. Therefore, as an alternative we identify doublets based on a combination of the inappropriate expression of cell-type marker genes coupled with elevated read counts. Marker genes that should be ubiquitously expressed (>90%) in one cell type (partition) and rarely expressed (<10%) in other cell types (partitions) were used to mark potential doublet cells. Cells with multiple inappropriate marker genes expressed (≥2) are tagged for removal. For example, *CD2* expression (cell adhesion molecule specific to T- and NKT-cells) was detected at higher than expected levels in VSMCs (3%) and ECs (2.1%), and thus these cells were excluded from analysis. An upward shift in read counts across all doublets relative to the population average was used to validate the doublet identification strategy. In addition, we perform gene co-expression network reconstruction using a custom partial correlations approach (described below), which is sensitive to rare outlying correlation events, and enables us to detect inappropriate co-expression of genes. Inappropriate networks reflective of unexpected cell-types were identified during the doublet detection process to validate our filtration strategy. Remaining cells were re-clustered producing six major partitions; macrophages, ECs, VSMCs, NKT cells, T- and B-lymphocytes (Fig. 2a-c). Patient-specific cells are shown in Extended Data Fig. 2b.

### Differential Gene Expression Analysis

Differential expression analysis was performed using Monocle^83^, which uses a generalized linear regression model approach, and was adjusted for patient specific and other confounder effects as covariates. Modeling includes an adjustment for patient identification as a corrective term. In order to have commensurate measures of differential expression, each gene’s expression level was normalized prior to fitting the model, allowing the resulting model coefficient to be interpreted directly as that gene’s effect size (normalized effect). The sign of the coefficient determines up-or down-regulation. A corrected p-value was computed for each coefficient using Benjamini and Hochberg. Consistency of gene expression differences at a biological process level was evaluated by Gene Set Enrichment Analysis applied to single-cell gene differential expression data ranked by normalized effect.

### Gene Networks

We reconstructed gene expression networks using a modified Weighted Gene Co-Expression Network Analysis^88^ approach we developed using partial correlations and applied to single-cell data. All pairwise gene-gene correlations are computed with partial correlations adjusted for the rest of the genome using a Penrose-Moore pseudo inverse with applied shrinkage parameter (uses the R package corpcor). The resulting matrix is linearized and then subjected to a false discovery analysis using R package fdrtool. Networks are reconstructed from the resulting top 20,000 most significant partial correlations by applying an FDR threshold to the pairwise co-expression edges (threshold estimated empirically from the distribution) with modules defined by Louvain clustering using default parameters^86^, where edge weights (distances) were set to the reciprocal of the absolute partial correlation. Because clustering was performed using weighted edges, FDR thresholds were relaxed and allowed to exceed 0.05 (mean FDR ~0.25).

### Gene Module Network Plotting and Selection of Significant Modules

Network plots were created using igraph’s built-in plotting functions. Cluster level plotting with colorization schemes aiding in visual interpretation of gene-to-gene relationships was used. For module plots containing a greater number of genes (over 100), additional filtering was used to help elucidate each module’s significant gene-to-gene relationships. For selected modules in the VSMC and EC cell types, module subnetworks were constructed by filtering and maintaining genes that were in the top 15 percent of those most connected within the network (higher ranked strength), along with all the modules’ differentially expressed genes. This resulted in modules on the order of 10 to 30 genes. Nodes were colorized gray for non-differentially expressed genes, and a cyan to red gradient was used for positive to negative normalized effect levels (qualified with q< 0.05). Dark red genes (nodes in the network) were indicative of genes significantly upregulated in AC cells, while dark cyan genes were lower expressing the AC cells.

In order to focus our analysis on key genetic drivers of disease processes, we chose to concentrate on selected modules exhibiting significant interactions between DE genes. This was determined by first setting a threshold of p<0.05 for differential expression overlap between modules (determined by one-tailed Fisher exact test). A contingency table was constructed for each module in the network based on the overlap of significant DE genes to the module genes. Modules do not overlap with each other due to the use of Louvain clustering. Modules with disproportionate numbers of DE genes rank as the most significant, while those with few or no DE genes rank as the least significant. This resulted in 8/31 modules being selected for further investigation in the VSMC dataset, and 7/36 modules in the EC dataset. Next, within each of these modules, genes were sorted by normalized effect and q-value<0.05 in order to better interrogate significantly differentially expressed genes. After this, the gene list was sorted by strength of connections, with >0.3 being used as a cutoff for genes to explore further, given the greater correlations with other genes observed with higher strength score. Modules with genes present satisfying these criteria moved forward with analysis. Within the chosen modules, genes were again sorted by normalized effect and q<0.05. Genes with NE >0.5 or <−0.5 were chosen and plotted, and those with signal strength >0.3 were chosen for further examination.

### Heat Maps

All heat maps were constructed from gene expression data, with individual cells plotted along the horizonal axis and genes plotted along the vertical axis. Prior to plotting, expression data was converted to binary form (on/off), with each gene plotted as on if the expression level of that gene had an RNA count of two or more. A binary distance method was then used to drive hierarchical clustering along both axes using complete-linkage clustering. All heat maps are accompanied by dot plots on the right side the heat map, showing gene expression levels for each of the cell subsets in the heat map. These dot plots are scaled based on the raw RNA counts.

### Extended Data

**Fig. 1:**
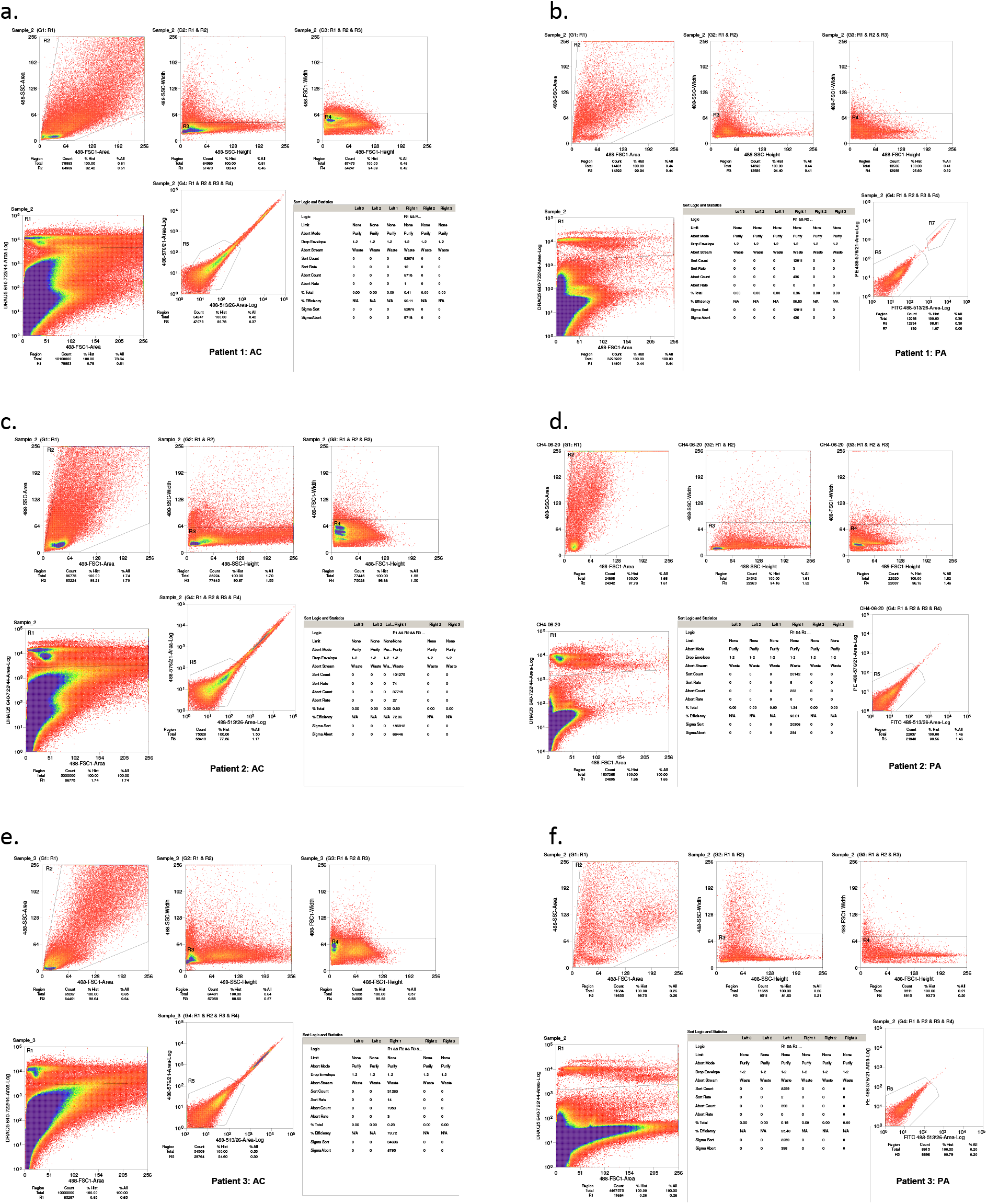
Fluorescence Activated Cell Sorting (FACS) of samples. **a-f,** Cells from dissociated carotid artery and plaque from both AC and PA samples were labeled with DRAQ5, a far-red excitation/ emission nuclear stain. Cells were distinguished from cellular debris by gating DRAQ5 positive events and doublets excluded using appropriate FSC and SSC gating.

**Fig. 2:**
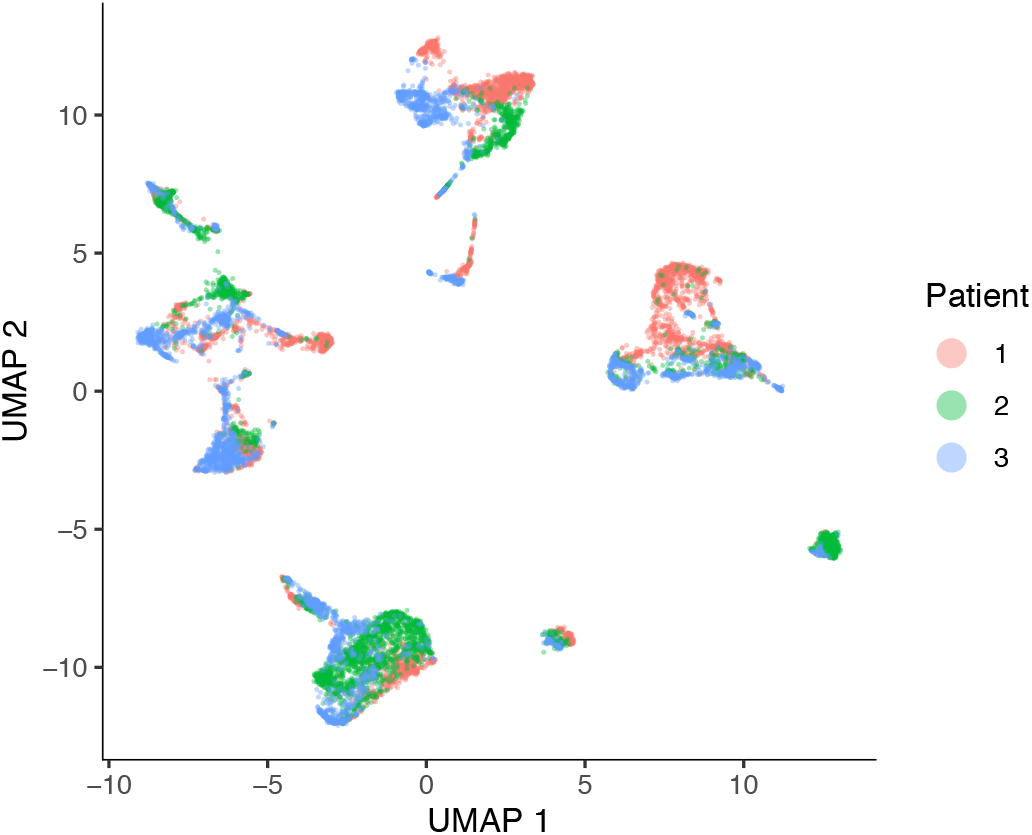
Aggregated patient data sets. **a,** UMAP of aggregated, down-sampled cell sets separated by patient.

**Fig. 3:**
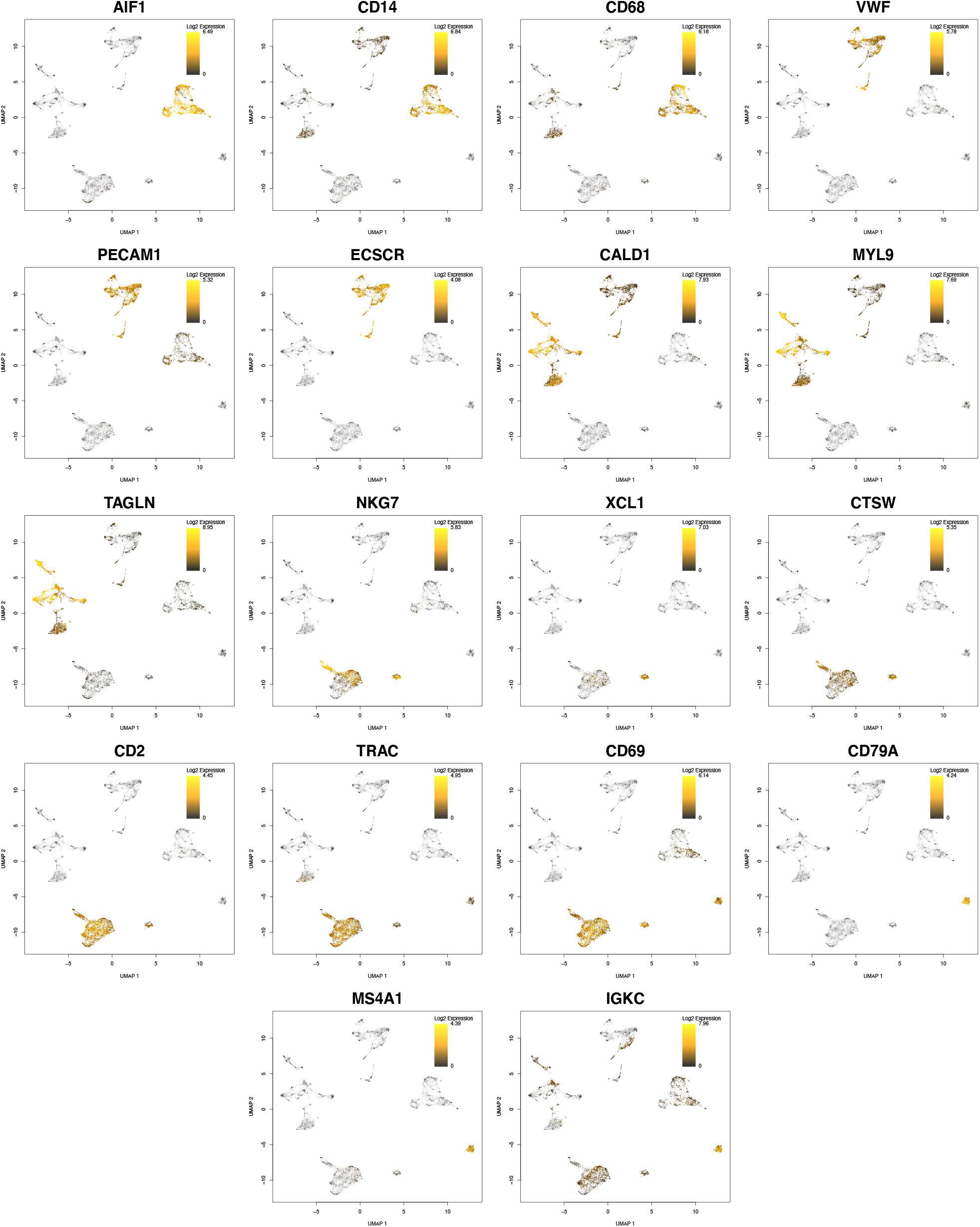
Gene markers for identified cell types. UMAP plots of selected cell-marker genes from dotplot (Fig. 2c). Color bar gradient indicates lowest (black) to highest (yellow) gene expression level.

**Fig. 4:**
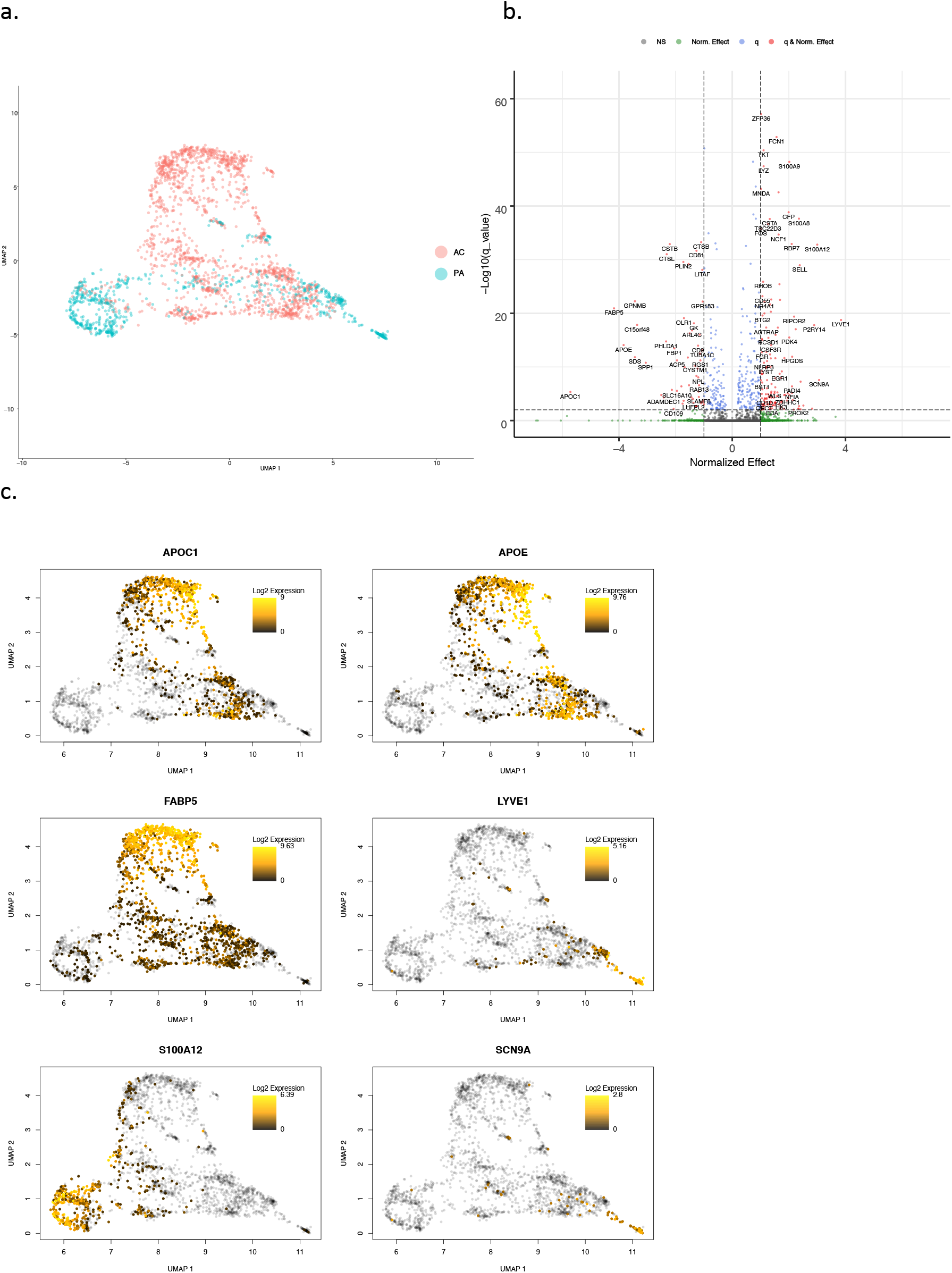
Differential expression analysis of macrophages. **a,** UMAP visualization of macrophages separated by anatomic location (n=2,456 cells). **b,** Volcano plots of the top differentially expressed genes in macrophages. Dotted lines represented q-value<0.01 and normalized effect >0.5 and <−0.5. **c,** UMAP plot of the top 3 AC and PA upregulated genes in macrophages. Color bar gradient indicates lowest (black) to highest (yellow) gene expression level.

**Fig. 5:**
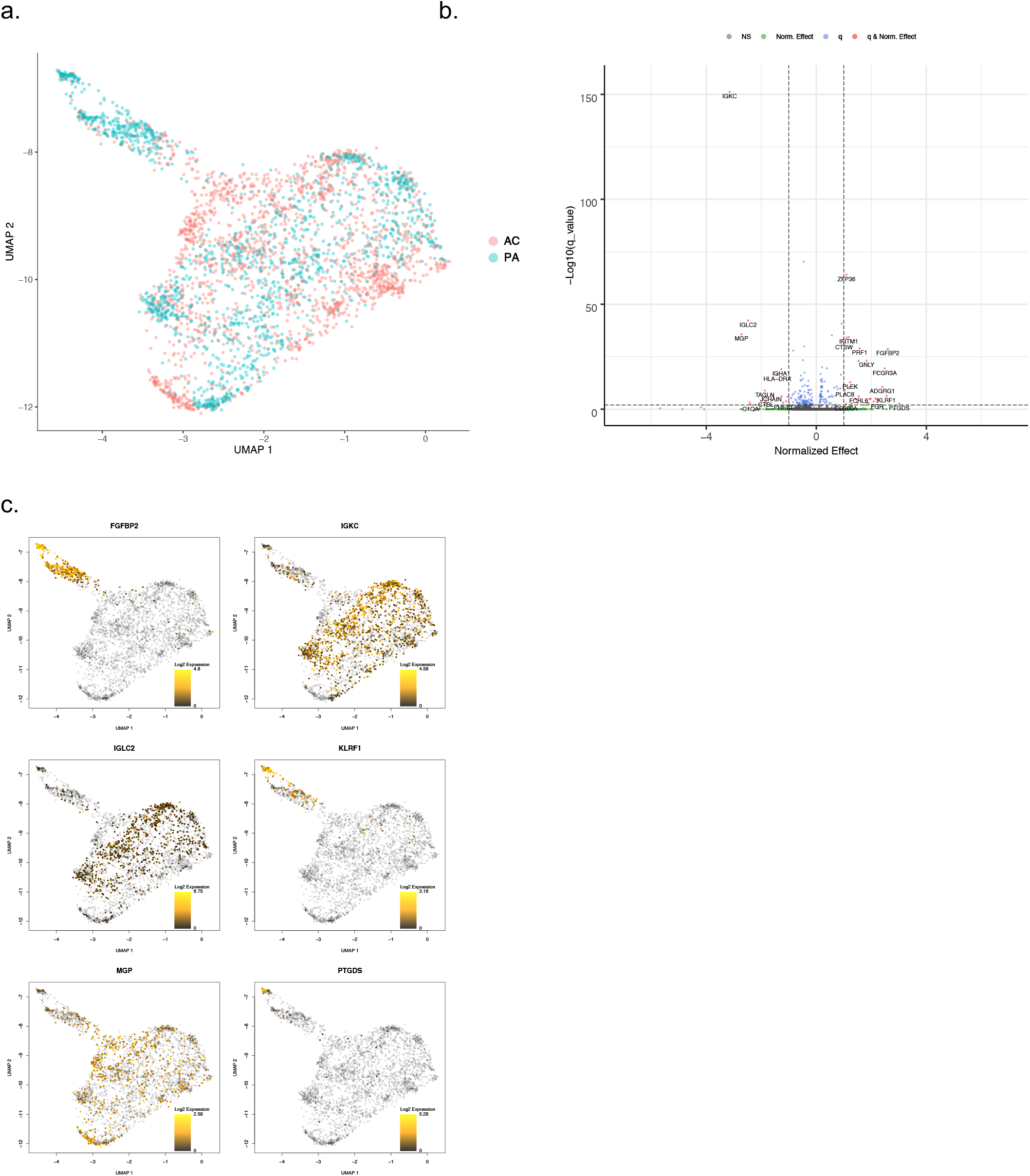
Differential expression analysis of T-lymphocytes. **a,** UMAP visualization of T-cells separated by anatomic location (n=3,402 cells). **b,** Volcano plots of the top differentially expressed genes in T-cells. Dotted lines represented q-value<0.01 and normalized effect >0.5 and <−0.5. **c,** UMAP plot of the top AC and PA differentially expression genes in T-cells. Color bar gradient indicates lowest (black) to highest (yellow) gene expression level.

**Fig. 6:**
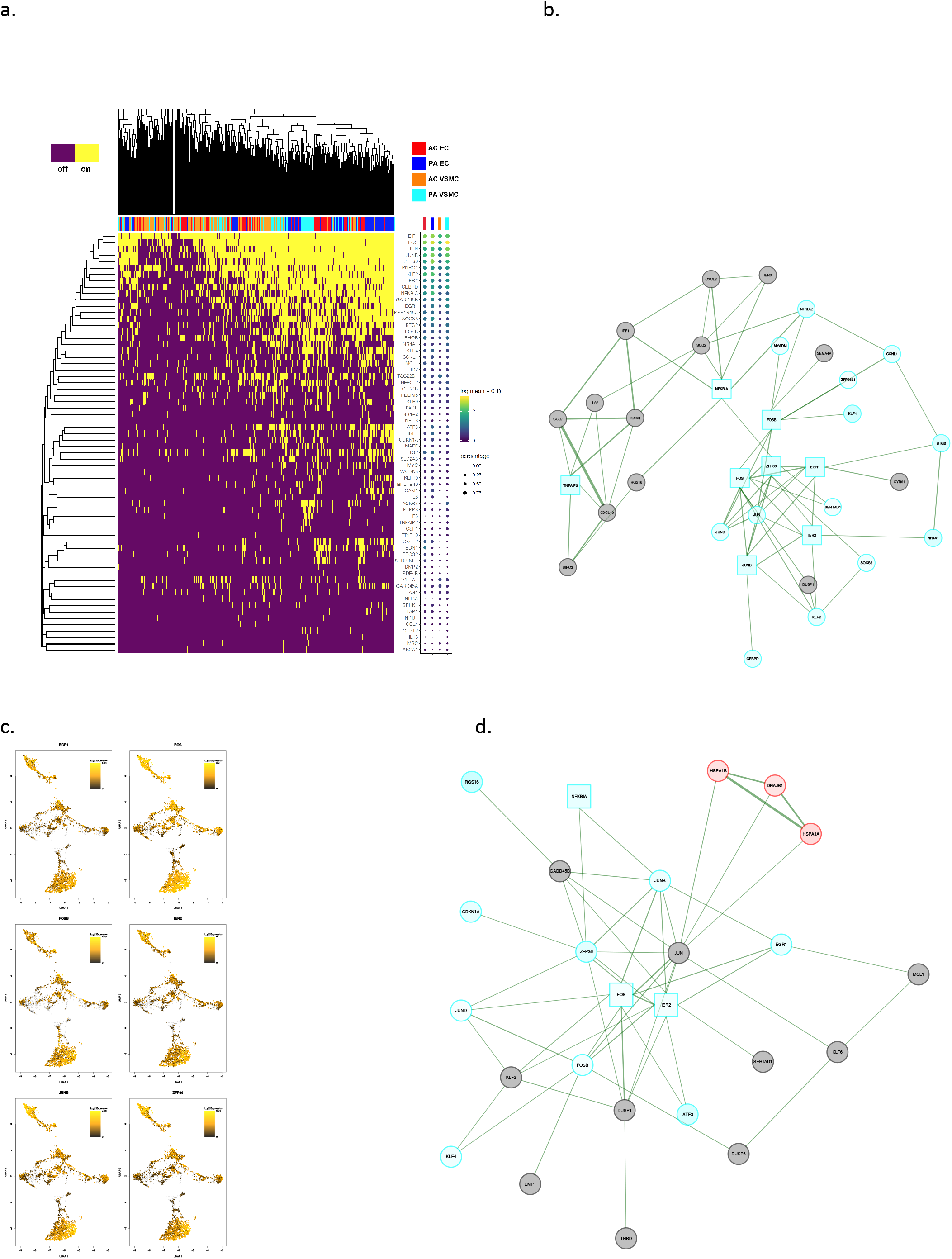
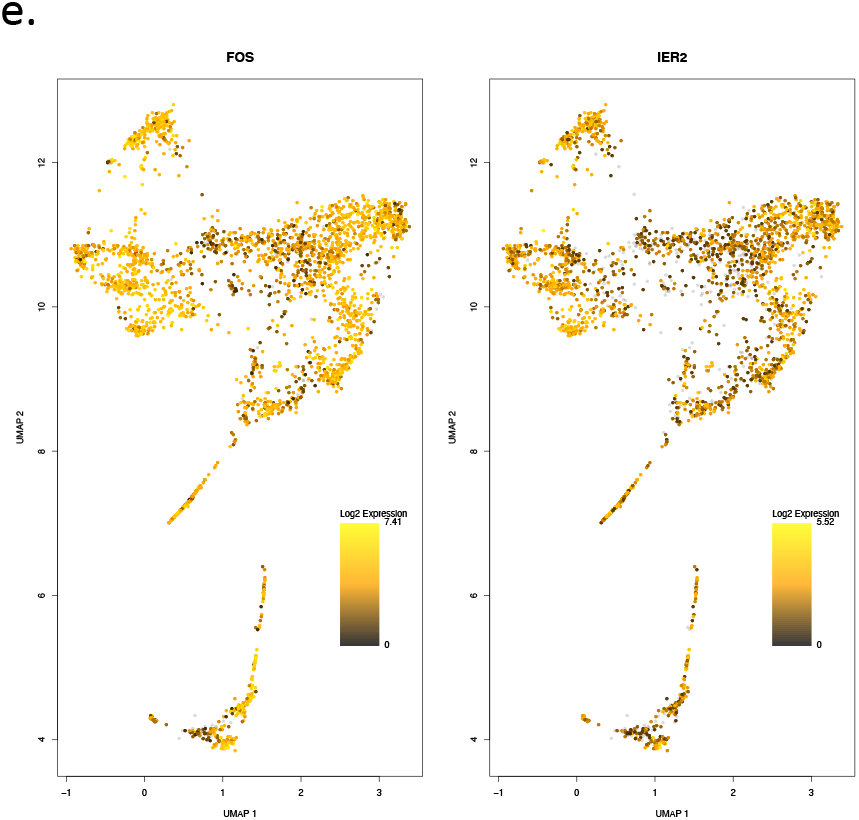
Gene set enrichment analysis and gene co-expression networks identify key gene drivers of TNFa signaling via NFkB hallmark biologic process. **a,** Fully clustered on/off heatmap visualization of overlap between significantly differentially expressed genes in VSMCs and ECs and leading edge EMT Hallmark genes generated by GSEA. Heatmaps are downsampled and represent 448 cells from each cell type and anatomic location (1792 total cells). A dotplot corresponding to gene expression levels for each cell type in the heatmap is included. Dot size depicts the fraction of cells expressing a gene. Dot color depicts the degree of expression of each gene. **b,d** Gene co-expression networks generated from VSMC Module 31 (b) and EC Module 36 (d) representing the TNFa signaling via NFkB hallmark from GSEA analysis. Genes are separated by anatomic location (red=AC genes, cyan=PA genes), differential expression (darker shade=higher DE, gray=non-significantly DEGs), correlation with other connected genes (green line=positive correlation, orange line=negative correlation) and strength of correlation (connecting line thickness). Significantly DEGs (q<0.05) with high connectivity scores (>0.3) are denoted by a box instead of a circle. **c,e** UMAP distribution of boxed genes from (b), (d), respectively. Color bar gradient indicates lowest (black) to highest (yellow) gene expression level. VSMCs=3,674 cells; ECs=2,764 cells.

**Fig. 7:**
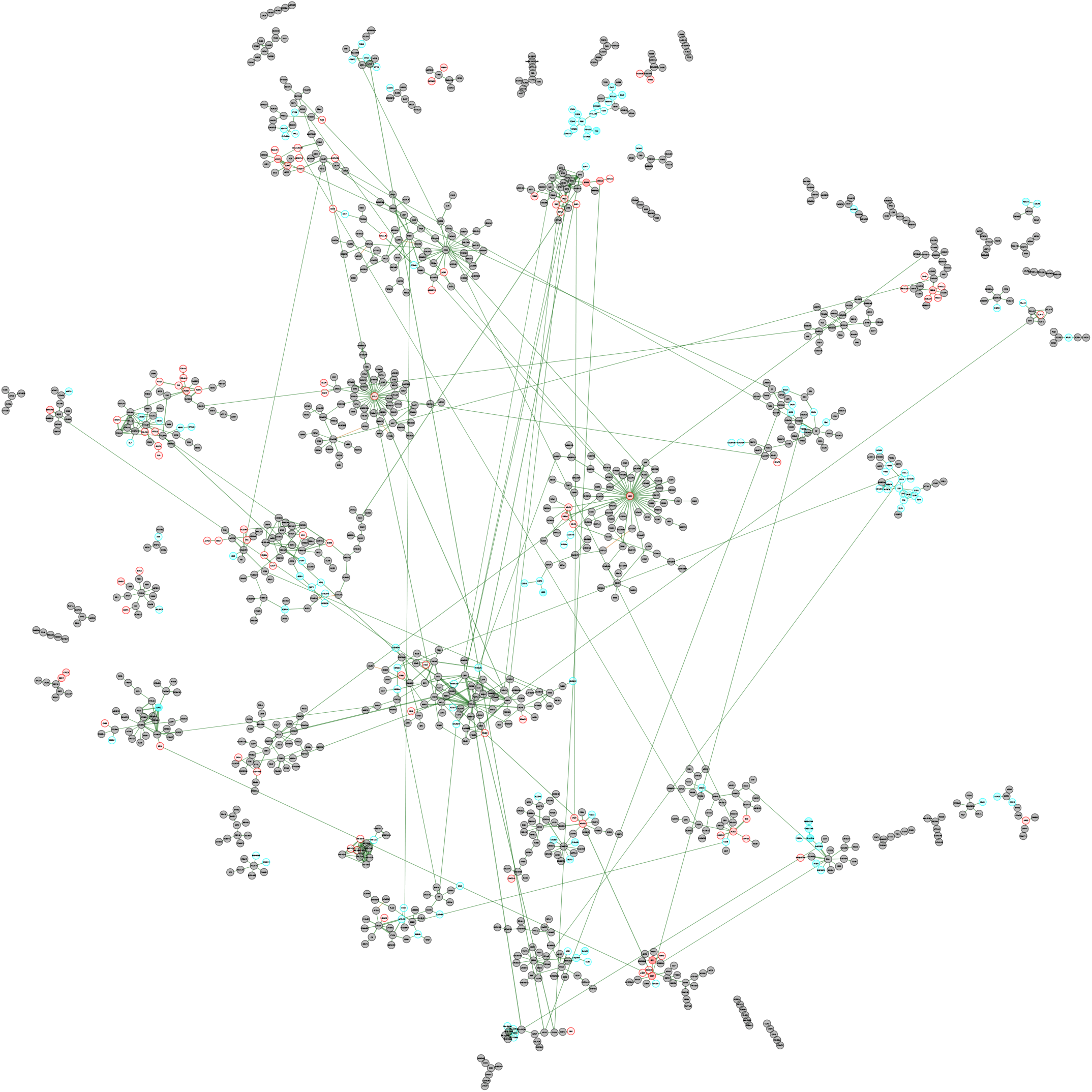
Full VSMC gene co-expression network. **a,** Gene co-expression network generated from all VSMC modules. Genes are separated by anatomic location (red=AC genes, cyan=PA genes), differential expression (darker shade=higher DE, gray=non-significantly DEGs), correlation with other connected genes (green line=positive correlation, orange line=negative correlation) and strength of correlation (connecting line thickness).

**Fig. 8:**
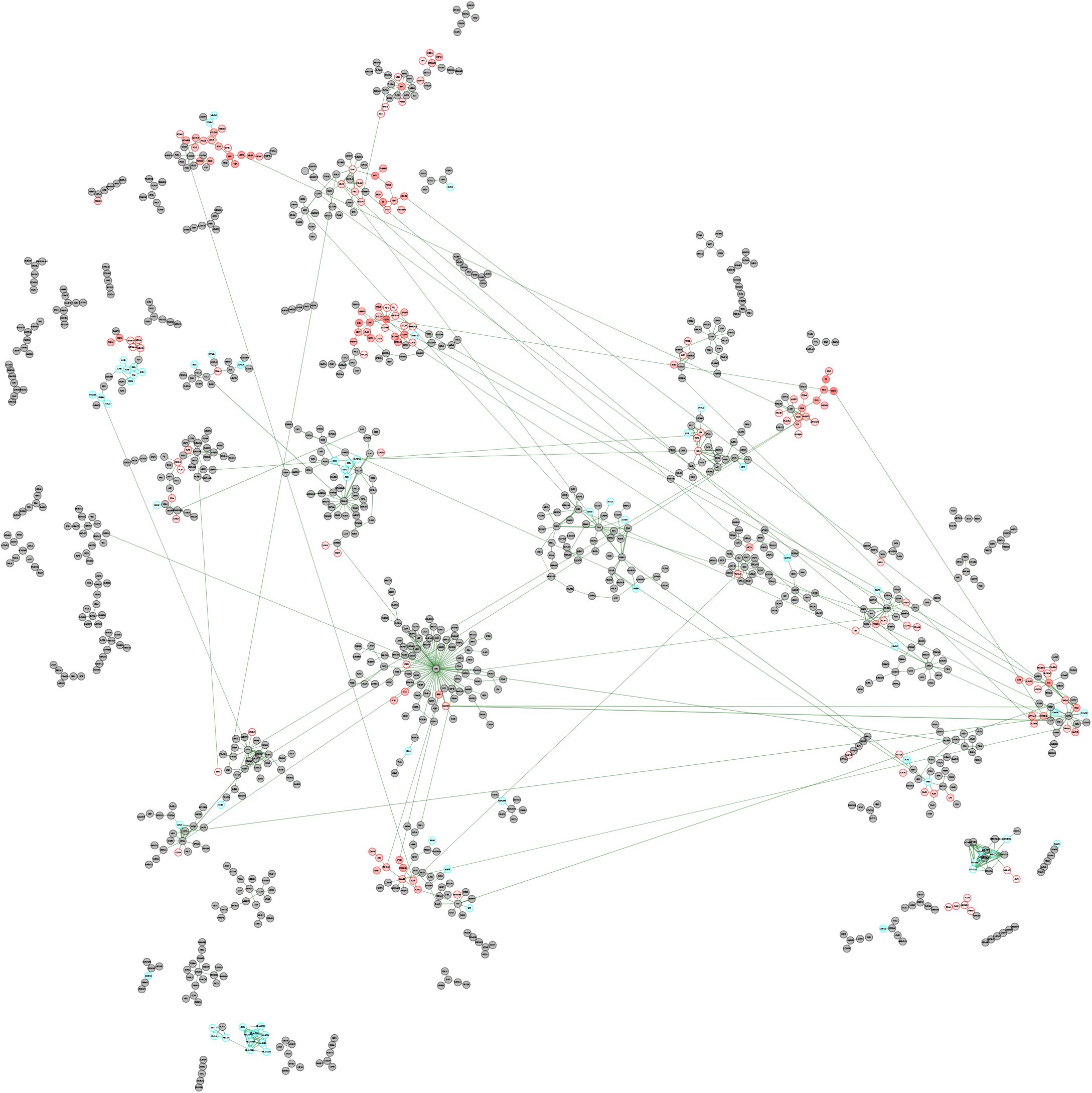
Full EC gene co-expression network. **a,** Gene co-expression network generated from all EC modules. Genes are separated by anatomic location (red=AC genes, cyan=PA genes), differential expression (darker shade=higher DE, gray=non-significantly DEGs), correlation with other connected genes (green line=positive correlation, orange line=negative correlation) and strength of correlation (connecting line thickness).

